# Lineage tracing identifies heterogeneous hepatoblast contribution to cell lineages and postembryonic organ growth dynamics

**DOI:** 10.1101/2023.01.18.524321

**Authors:** Iris. A. Unterweger, Julie Klepstad, Edouard Hannezo, Pia R. Lundegaard, Ala Trusina, Elke A. Ober

## Abstract

To meet the physiological demands of the body, organs need to establish a functional tissue architecture and adequate size as the embryo develops to adulthood. In the liver, uni- and bipotent progenitor differentiation into hepatocytes and biliary epithelial cells (BECs), and their relative proportions comprise the functional architecture. Yet, the contribution of individual liver progenitors at the organ level to both fates, and their specific proportion is unresolved. Combining mathematical modelling with organ-wide, multispectral FRaeppli-NLS lineage tracing in zebrafish, we demonstrate that a precise BEC to hepatocyte ratio is established (i) fast, (ii) solely by heterogeneous lineage decisions from uni- and bipotent progenitors, and (iii) independent of subsequent cell type-specific proliferation. Extending lineage tracing to adulthood determined that embryonic cells undergo spatially heterogeneous three-dimensional growth associated with distinct environments. Strikingly, giant clusters comprising almost half a ventral lobe suggest lobe-specific dominant-like growth behaviours. We show substantial hepatocyte polyploidy in juveniles representing another hallmark of postembryonic liver growth. Our findings uncover heterogeneous progenitor contributions to tissue architecture-defining cell type proportions and postembryonic organ growth as key mechanisms forming the adult liver.

## Introduction

Liver formation requires the timely differentiation of multipotent progenitor cells into specific cell types that form the building blocks of the organ. Relative proportions of these cell types are critical for establishing a specialized tissue architecture mediating physiologic liver functions. During embryonic and postembryonic growth, the liver increases in size to meet the growing physiological demands. Yet, how individual progenitors contribute to distinct cell lineages and subsequent growth are fundamental questions in organogenesis.

The liver consists mostly of hepatocytes and biliary epithelial cells (BECs), also called cholangiocytes, which together with mesenchymal cell types, are arranged in a characteristic architecture executing essential liver functions. On the tissue scale, the mammalian liver is divided into liver lobules with a central vein and portal triads at the edges, consisting of portal veins, arteries and biliary ducts, whereas hepatocytes distribute within the lobule along sinusoids connecting the main vessels [1]. Zebrafish lack portal triads, instead, the central vein resides in the core of each lobe, and the portal veins at the periphery [2,3]. Hepatocytes align along sinusoids between the two veins. Both sinusoids and the intrahepatic bile ductules are organised throughout the lobe in complementary mesh-like networks [4].

During zebrafish development, hepatic progenitors, called hepatoblasts, are specified in the ventral foregut endoderm by signals from the adjacent mesoderm around 23 hours post fertilization (hpf) [5–7], and migrate to form the nascent liver bud [8]. Immunohistochemistry studies of the rat liver initially suggested the bipotent nature of hepatoblasts, to differentiate into BECs or hepatocytes [9]. Bipotency was subsequently demonstrated *in vitro* by culturing mouse hepatoblasts isolated by selected surface markers in respective culture media [10,11] and more recently in organoids [12]. Lineage tracing of early definitive foregut endoderm in mice, labelled at E7.75, showed contribution to both lineages pointing to bipotentiality. However, recombination was induced prior to liver specification [13]. Instead, tracing of Lgr5^+^ hepatoblasts from E9.5, representing 2% of hepatoblasts at this timepoint, showed the functionally heterogeneous contribution of uni- and bipotent hepatoblasts to hepatocytes and BECs when focussing on the portal triad [12]. Therefore, comprehensive information about hepatoblast potential at the population level and their contribution across the entire lobe is missing. The identified unipotent hepatoblasts exclusively contributed to the hepatocyte but not the BEC lineage [12]. Whether heterogeneous hepatoblast potential represents a conserved strategy in vertebrate liver formation is an open question. Furthermore, it is unknown how the precise cell type proportions critical for a functional organ architecture are established.

Transcriptional profiling in mice suggests that the transition from hepatoblasts to hepatocytes occurs by default in the absence of specific inductive signals, whereas the hepatoblast to BEC transition represents a regulated process [14,15]. Notch signalling plays a recurrent and essential role during BEC differentiation and BEC duct formation across species [16–19]. In the zebrafish liver, Notch ligands regulate BEC specification [18] and the Notch reporter *tp1:EGFP* is expressed in BECs first at 45 hpf [16]. In parallel, hepatocyte differentiation begins between 40 and 60 hpf [4].

Once the nascent tissue organisation is established, the liver transitions into a growth phase [19]. This occurs in zebrafish around 5 days post fertilization (dpf), when the liver consists of two lobes, the left and right lobe, and takes up organ-specific functions [20]. As the liver enlarges during postembryonic growth, a third liver lobe, the ventral lobe, arises [21]. While the molecular mechanisms of liver cell type differentiation are gradually being elucidated [4], postnatal growth and the transition to the adult organ remain generally poorly understood [22,23]. To accommodate the 900-fold increase in cell number between 5 dpf and 1.5 years, each embryonic liver cell in zebrafish divides theoretically ten times [24]. Lineage tracing over a similar period revealed that new hepatocytes arise exclusively from the proliferation of existing hepatocytes [24]. However, BECs can also transdifferentiate into hepatocytes and contribute to the hepatocyte pool in homeostasis [25]. Similar differences have been seen in mice, attributed mostly to diverse lineage tracing approaches [26]. Furthermore, the contribution of individual hepatoblasts to the growth of the adult liver remains unknown: for example, do all hepatoblasts produce an equal number of progeny or do some generate more than others [27,28]? Mechanistically, this could be controlled spatially, for instance, by growth zones at the organ periphery during development [29–31] or regionally within the lobule [32]. Overall, there is a large gap in our understanding of postembryonic liver growth across all liver lobes, as well as between species.

Combining lineage tracing with whole-mount imaging and mathematical modelling in zebrafish, we here show that solely heterogeneous lineage contributions of progenitors are sufficient for establishing the precise proportion of BECs and hepatocytes comprising the functional liver architecture. Furthermore, by morphological and clonal studies, we demonstrate that embryonic cells contribute heterogeneously to postembryonic liver growth, including giant clusters driving the distinct growth behaviour of the ventral lobe during metamorphosis.

## Results

### Mathematical modelling for establishing a precise BEC to hepatocyte proportion

To elucidate how the correct number and proportion of hepatocytes and BECs arise in liver development (Fig. 1A), we first determined their cell numbers at 5 dpf when hepatic tissue organisation is established, and the liver takes up function (Fig. 1B). Visualizing hepatocytes and BECs by expression of Hnf4α [33] and the Notch reporter *tp1:EGFP* [16], respectively, we showed that hepatocytes outnumber BECs by 9-fold (Fig. 1C). Asking how this 1:9 BEC to hepatocyte ratio is established, we turned to mathematical modelling considering two parameters: lineage potential of the initial hepatoblast population and proliferation rates of the differentiated cell types (Figs. 1D – F, S1A-C). The simplest approach to achieve this 1:9 ratio would solely comprise unipotent hepatoblasts, of which 10% contribute to the BEC lineage and 90% generate hepatocytes. However, since mammalian liver formation involves bipotent hepatoblasts [12], they were included in all our models.

**Fig. 1.**
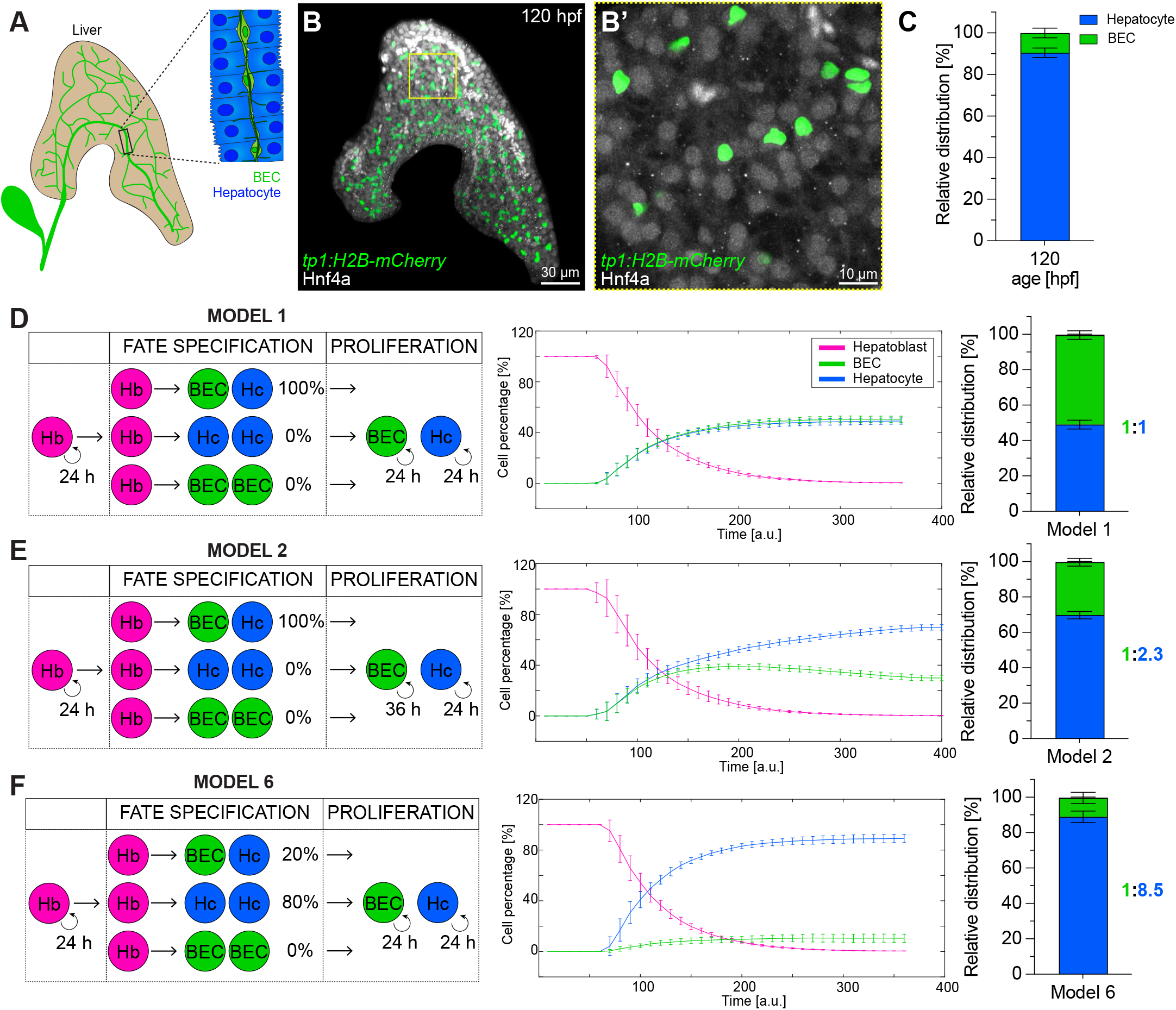
Establishment of BEC and hepatocyte lineages: *in vivo* cell type quantification and *in silico* modelling. (A) Schematic of 5 dpf liver, highlighting the biliary network. (B-B’) Maximum projection (200 µm z-stack) of a 120 hpf liver expressing *tp1:H2B-mCherry* (BEC) and stained for Hnf4ɑ (hepatocyte). (N=4, n≥12) (C) Relative distribution of BECs and hepatocytes at 120 hpf (N=4, n≥12). (D-F) Mathematical models simulating hepatoblast differentiation employing different parameter combinations: proliferation rates of differentiated cell types is equal (D,F) or slower in BECs (E); Hepatoblasts are either all bipotent (D,E) or represent a heterogeneous population with mixed probabilities for uni-or bipotent differentiation (F). Plots showing the simulated cell proportions over simulation time (n=10) and the final cell type ratio in bar graphs. Hb = hepatoblast, Hc = hepatocyte.

First, given that a 1:1 BEC to hepatocyte ratio is produced when all hepatoblasts are equipotent (model 1; Fig. 1D), we decreased in model 2 the division rate of BECs by an estimated 50% generating a 1:2.3 BEC to hepatocyte ratio, suggesting that moderately different cell division times alone are not sufficient to produce the *in vivo* proportion (model 2; Fig. 1E). Next, varying the composition of hepatoblasts by introducing uni- and bipotent potentials lead to ratios with a higher hepatocyte fraction (model 3-6, Fig. 1F, S1A-C). The model producing an outcome (ratio 1:8.5) closest to the 1:9 *in vivo* ratio contains 80% unipotent hepatocyte-producing hepatoblasts and 20% bipotent ones, exhibiting equal proliferation rates (model 6; Fig. 1F). Adding a low probability of 5% unipotent hepatoblasts contributing solely BECs, resulted in a 1:5.7 (model 4, Fig. S1B) compared to the 1:8.5 ratio in model 6 (Fig. 1F). Whereas, decreasing BEC proliferation rates in this model similar to above, established a 1:11.8 ratio over time (model 5; Fig. S1C).

It is noteworthy that in the various models respective BEC to hepatocyte ratios are established with different velocities. Specifically, a BEC-hepatocyte equilibrium is reached faster in models 4 and 6 with the greatest hepatoblast heterogeneity, while solely altering proliferation rates between cell types in model 2, takes about 3-times longer.

In summary, mathematical modelling predicts that the *in vivo* 1:9 BEC to hepatocyte ratio cannot be established by decreasing the BEC proliferation rate alone. In contrast, a heterogeneous progenitor lineage potential is sufficient to achieve such proportions.

### BEC and hepatocyte proliferation dynamics during embryonic development

Based on mathematical models 2 and 5, unequal proliferation rates can contribute to a differential BEC to hepatocyte ratio. To test whether *in vivo* division rates between BECs and hepatocytes are similar or differ, we examined cell proliferation by 5-ethynyl-2’-deoxyuridine (EdU) incorporation between 48-144 hpf. Starting at 48 hpf, 16-18% of both cell types proliferate, at a rate continuously decreasing until 120 hpf, when only 2-4% are EdU-positive (Fig. 2A). Both the total number of BECs and hepatocytes increases by 8.4-fold between 48 and 120 hpf (Figs. 2B,C), while the liver volume increases disproportionately by 20-fold within the same timeframe (Fig. 2D).

**Fig. 2.**
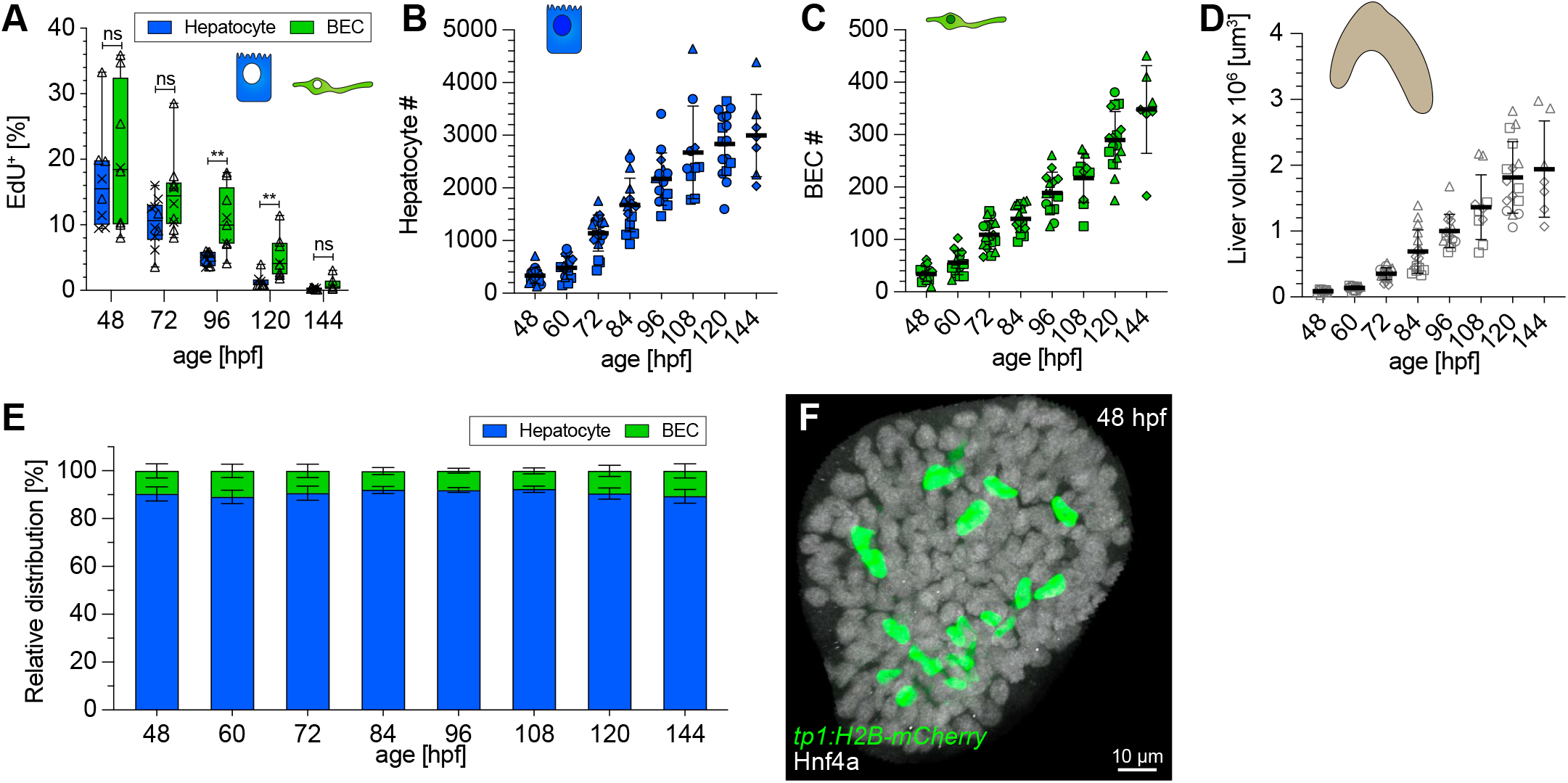
Hepatic proliferation dynamics and early establishment of a 1:9 BEC:hepatocyte ratio during embryonic development. (A) The proportion of EdU^+^ proliferating hepatocytes and BECs steadily decreases over time (N=2, n≥8). (B-C) Graph showing hepatocyte (B) and BEC (C) cell numbers during development (N=4, n≥12). (D) Quantification of total liver volume during development determined in embryos in BABB (N=4, n≥12). (E) Relative distribution of BECs and hepatocytes during development from 48 to 144 hpf (N=4, n≥12). (F) Maximum projection (20 µm z-stacks) of a 48 hpf liver expressing *tp1:H2B-mCherry* (BEC) and stained for Hnf4ɑ (hepatocyte).

Our results reveal minimal differences between BEC and hepatocyte proliferation rates, suggesting progenitor potential as a major factor for the establishment of the 1:9 BEC to hepatocyte ratio *in vivo*, mirrored in mathematical models 4 and 6 (Figs. 1F, S1B).

### The mature BEC to hepatocyte ratio is already established early in liver development

Another distinguishing hallmark between the different models is the velocity, up to 3-fold different, by which the BEC to hepatocyte ratio arises. We determined cell type numbers throughout development to assess when the 1:9 BEC to hepatocyte ratio is established after cell type specification *in vivo*. Unexpectedly, the 1:9 ratio of BECs to hepatocytes is already reached at 48 hpf (Fig. 2E) only a few hours after the onset of BEC differentiation (Fig. 2F). This finding together with similar BEC and hepatocyte proliferation rates (Fig. 2A) strongly supports models 4 and 6, in which hepatoblast heterogeneity is sufficient to rapidly establish the final ratio and maintain it over time.

Asking whether this distinct cell type ratio is characteristic for the differentiating embryonic liver or important for tissue functionality, we next examined *in vivo* cell numbers in postembryonic livers (Figs. S1D-G). We determined a 1:7.9 BEC to hepatocyte ratio in juvenile and an average 1:8.75 in adult livers (periphery: 1:6.4, and center:1:11.1), suggesting that similar cell type proportions are maintained from the embryonic to the mature liver.

### Lineage tracing identifies uni- and bipotent hepatoblasts *in vivo*

Next, to investigate hepatoblast potential, we applied unbiased lineage tracing strategies. Thereby, also addressing the fundamental question as to whether hepatoblast potential is conserved strategy across species. To this end, we used the multicolour labelling system FRaeppli-NLS, which upon conditional activation, stochastically labels nuclei with one of four fluorescent proteins (FPs): TagBFP, mTFP1, E2-Orange, or mKate2 [34] (Fig. 3A). The spectra of the FRaeppli FPs allow combination with transgenic *tp1:EGFP* expressed in BECs. To ensure that the *fraeppli-nls* transgene is expressed in progenitors and maintained in hepatocytes and BECs, we used *prox1a:kalTA4* as a driver (Fig. 3A), given Prox1 is expressed in hepatoblasts and differentiated hepatocytes and BECs (Fig.S2A) [18]

**Fig. 3.**
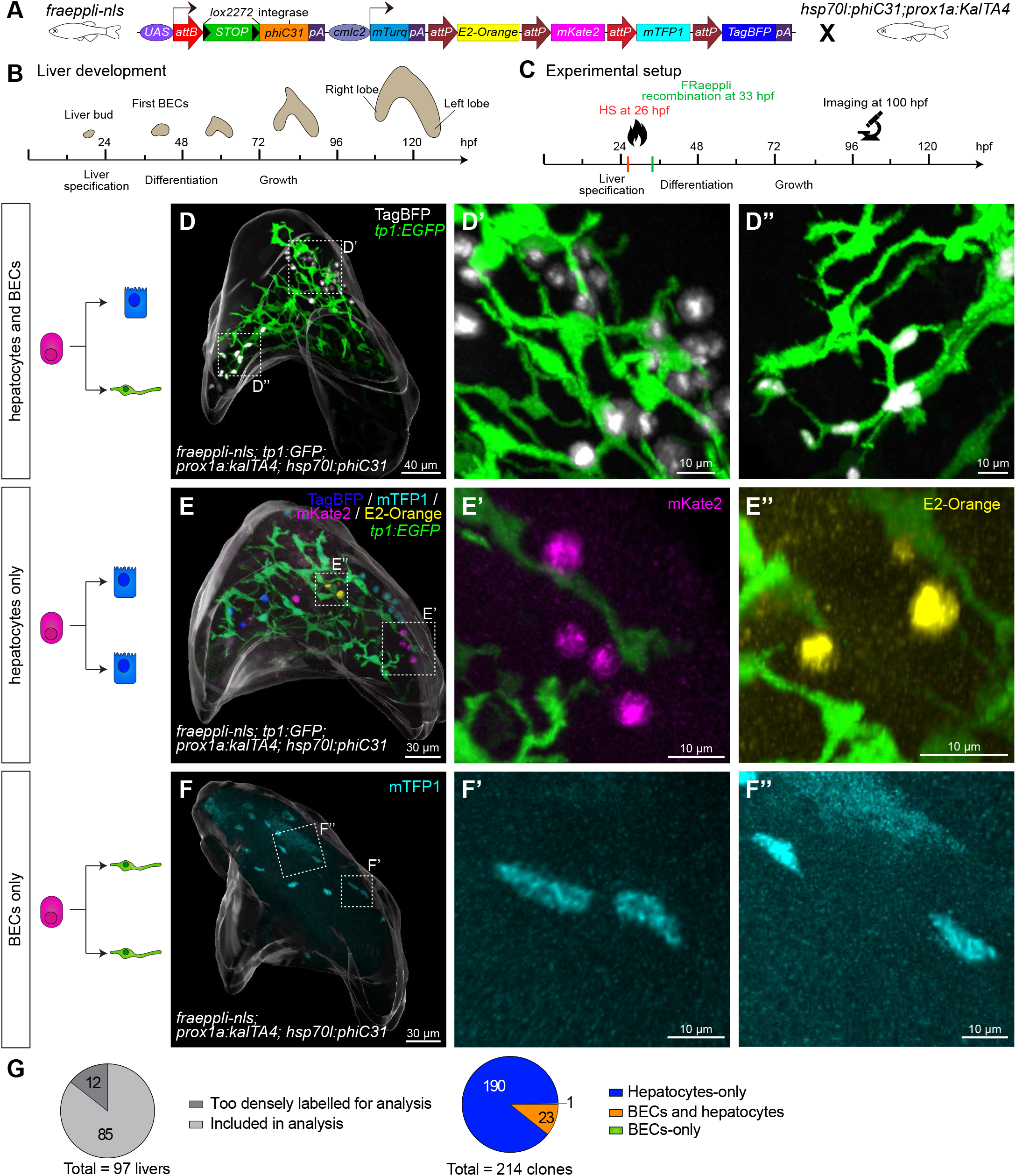
Quantitative lineage tracing identifies uni- and bipotent hepatoblast contributions during lineage decisions. (A) Schematic of FRaeppli-NLS cassette including attB and attP sites for PhiC31-mediated recombination and the four FRaeppli fluorescent proteins: TagBFP, mTFP1, mKate2 and E2-Orange. Recombination is induced by combining *fraeppli-nls* with *hsp70l:phiC31; prox1a:kalTA4*. (B) Key steps of liver development in zebrafish: after hepatoblast specification, the differentiation into BECs and hepatocytes is initiated at around 42 hpf. Differentiated cells acquire polarity and form a functional architecture by 120 hpf. (C) Experimental strategy for tracing progeny of individual hepatoblasts using *fraeppli-nls*: Heat shock at 26 hpf controls PhiC31 expression followed by attB-attP recombination. Embryos were fixed at 100 hpf for analysis. (D-F) Whole-mount livers at 100 hpf showing: (D) mixed clone composed of hepatocytes (D’) and BECs (D’’) (N= 6, n=23); (E) clones formed by hepatocytes only (E’-E’’) (N= 6, n=190) and (F) BEC-only clone in mTFP1 (F’-F’’) (N= 6, n=1). (D-F) An overall segmentation of the whole liver tissue is shown in transparent grey. (G) Pie charts showing the total number of labelled embryos and clones with manually assigned lineage contributions (N=6, n=214; in two of the six experiments nuclear shape indicated BEC fate).

To label cells at the hepatoblast stage, prior to cell differentiation, we combined *fraeppli-nls* with *hsp70l:phiC31* and induced recombination by heat-shock at 26 hpf (Figs. 3B-C). PhiC31 maturation and attB/attP-mediated recombination take about 5-9 hours [34] and thus would be completed prior to fate commitment (Figs. 3B-C, S2B). Based on total liver cell numbers and cell doubling times, we estimated that FRaeppli-labelled hepatoblasts would undergo maximally one cell division before lineage decision at around 40-42 hpf (Fig. S2C). For analysis, we fixed embryos at 100 hpf, when all four FRaeppli FPs are strongly expressed (Fig. S2D-E). To determine cell fates within a clone, we combined *fraeppli-nls; prox1a:kalTA4; hsp70l:phiC31* with *tp1:EGFP* to distinguish BEC from hepatocyte fate. In addition, samples without *tp1:EGFP* expression were included using nuclear shape as an indicator of cell fate, given that at 100 hpf BEC nuclei are mostly elongated, while hepatocyte nuclei are round (Fig. S2A) [35]. Given that cell rearrangement can lead to fragmentation and merging of clones [34,36], we established rigorous rules to define clones. Considering both the range of cell movement determined by *live*-imaging and the volume increase of the entire organ (Fig. S2C,F), clones were defined as labelled cells of the same colour located within a 70 µm radius (Fig. S2G).

With this strategy, 214 clones were analysed for their lineage contribution to hepatocytes and BECs. A recombined clone was found in 1.8% of the control samples, indicating low unspecific recombination. Interestingly, out of all heat-shock induced embryos, 78% exclusively labelled hepatocytes, while the remaining 22% displayed clones also containing labelled BECs. Overall, we observed three distinct clonal outcomes: first, mixed clones accounting for 10.7% of cases and indicated to originate from bipotent hepatoblasts (Fig. 3D). Second, the majority of clones, 88.8%, consisted of hepatocytes only, in line with a unipotent hepatoblast potential (Fig. 3E). Finally, in one liver, representing 0.5% of clones, all labelled cells were BECs suggesting a unipotent hepatoblast with a BEC-restricted potential. Notably, in this pure BEC clone, cell fate was assigned by nuclear shape (Fig. 3F). Lineage tracing of stochastically labelled hepatoblasts reveals their heterogeneous potential, encompassing bipotent hepatoblasts, as well as a high fraction of unipotent hepatoblasts contributing predominantly to the hepatocyte lineage (Fig. 3G), as predicted by mathematical model 6 (Fig. 1F).

### Hepatoblasts contribute stochastically to embryonic liver growth

Hepatoblast lineage tracing revealed a striking range of clone sizes, raising the question as to whether their proliferative capacity and contribution to the overall liver is controlled at the level of the individual progenitor or stochastic. We focused on hepatocyte clones because BEC-containing clones frequently expand into a widely ramified network; therefore, size determination is challenging. Clone size varied, from 1-33 cells (Fig. 4A,C-E), independent of clone colour (Fig. 4A). Given the range in clone size, we converted each cell number into the number of cell divisions each progenitor had undergone. This showed that the largest clone arose from a single progenitor that underwent six divisions, the majority, 22.1%, divided once, while a substantial number of hepatoblasts, 7.9 %, did not divide (Fig. 4B). Comparing this clonal division range to a Poisson distribution, which is the simplest null model where the distribution arises simply from the stochasticity of division timing of a single homogeneous population. The good fit between data and the simple model, in the absence of any free parameter apart from the average division time, suggests a fully stochastic proliferation behaviour (Figs. S3A,B).

**Fig. 4.**
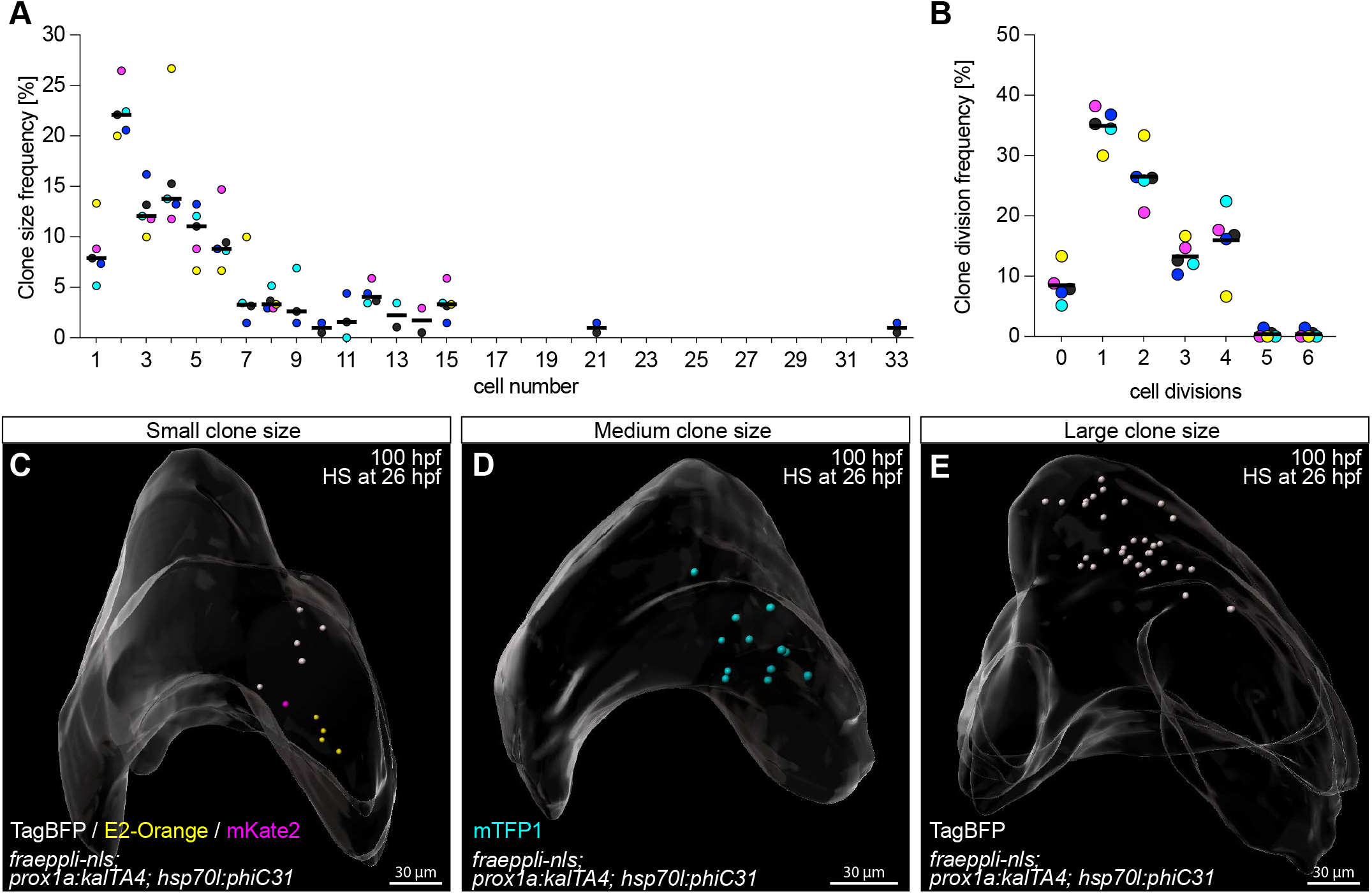
Quantitative lineage tracing of hepatoblasts during embryonic development identifies heterogeneous growth behaviour. (A) Frequency of manually-assigned clone sizes. (B) Distribution of the corresponding number of cell divisions for each clone. Clone colours are plotted in blue (TagBFP), turquoise (mTFP1), magenta (mKate2) and orange (E2-Orange); the mean of all colours is represented in black. (C) Whole-mount of 100 hpf liver showing several clones, including a mKate2^+^ 1-cell clone (N=6, n=15). (D) Liver with a medium size 12-cell mTFP1^+^ clone (N=6, n=7). (E) Whole-mount of a 100 hpf liver with a large 33-cell TagBFP^+^ clone (N=6, n=1). (C-E) Labelled cells are represented as segmented nuclei and an overall segmentation of the whole liver tissue is shown in transparent grey.

To rule out a bias arising from manual clone assignment, we defined clonality mathematically considering all labelled cells per liver (Fig. S3C). We first calculated the probability that a given cell had a neighbour with the same or a different colour (Fig S3D) [37,38]. Then, looking at subsets of livers with a defined number of total labelled cells, revealed that clonality assignment becomes unprecise when a threshold of 40-50 labelled cells per liver is exceeded (Fig. S3D). This also determined that clones of different colours are usually at least 50 µm apart, suggesting that labelled cells more than 50 µm apart should not be considered clones. Next, we compared the size distribution of manually defined clones to theoretically reconstructed clones (Fig. S3E). This analysis revealed that including samples with less than 40 labelled cells and grouping cells within a 45 µm distance between clones represents a suitable ruleset. Interestingly, the reconstructed clone size distribution was well fitted by a single exponential distribution, which is the theoretical expectation for a population undergoing stochastic division as inferred from the distribution of division numbers above, and thereby confirmed that the manual ruleset was a valid approximation (Fig. S3E). It is noteworthy that the average clone size of 4.5 cells (2.1x clone division rate) resulting from manual clone assignment (Fig. 4A) did not match the expected average of 16-cell clones (4x clone division rate) based on the total liver cell numbers between the time of recombination and analysis (Fig. S3F) [39], suggesting that likely larger clones were missed based on our strict, experimentally derived rule set. In summary, the unbiased mathematical approach confirmed that the manually defined clones and the resulting cell numbers are not or only minimally influenced by tissue rearrangements *in vivo*.

### Embryonic cells contribute heterogeneously to postembryonic growth

Next, we asked how constituent progenitors contribute to postembryonic organ growth as the liver dramatically increases in size and changes shape, including the *de novo* formation of a third liver lobe (Fig. 5A). Long-term lineage tracing experiments were performed to investigate whether the growth contribution of individual hepatoblasts is uniform throughout the organ, or whether some contribute minimally while others greatly. Similar to the lineage tracing experiments during development, we employed the FRaeppli-NLS system in combination with *hsp70l:phiC31*; *prox1a:kalTA4* to induce labelling of hepatoblasts by heat shock at 26 hpf (Figs. 5B), followed by qualitative analysis of the spatial patterns clusters exhibit in postembryonic livers. For that, we acquired 3D-data sets of 79 adult *fraeppli-nls; prox1a:kalTA4; hsp70l:phiC31* livers. In these recombined livers, a group of labelled cells is termed ‘cluster’, since we cannot exclude that cells labelled in the same colour are the progeny of more than one hepatoblast. First, 11.4% of recombined livers displayed clusters that distribute along the central veins in the core of the liver lobe (Figs. 5C-C’). In most of these cases, clusters are oriented along the anterior-posterior axis of the fish. Second, 30% of recombined livers clusters distribute in a stripe-like fashion, consistent with cells proliferating and arranging along endothelial sinusoids (Fig. S4G), creating parallel interspaced stripes. Specifically, in the lobe core, proximal-distal stripe-like clusters are oriented perpendicular to the central vein, such that labelled cells extend from the central vein to the margin of the lobe (Fig. 5D-D’). Similarly, striped clusters were also present in the anterior part of the liver, which connects the three lobes (Fig. S4A-A’). Finally, 3.8% of recombined samples exhibited some unexpectedly large clusters, which we termed ‘giant clusters’, since they occupied nearly half a lobe (Fig.4F-G’, S4B-C). Remarkably, giant clusters comprised 6-11% of the total liver volume, in contrast to an expected 0.7% assuming that all hepatoblasts at the timepoint of labelling proliferate equally. In contrast to the other two cluster shapes, which can be found throughout all three lobes, giant clusters were located exclusively at the tip of the ventral lobe and extended towards the left lobe (Figs. 5F-G). For one giant cluster, we traced the embryonic origin to a single mOrange2-labelled cell at the periphery of the left lobe at 5 dpf (Figs. 5E-F). Importantly, a TagBFP clone in the same sample, forms a much smaller clone in the adult liver in the anterior ventral lobe (Fig.5E-F), suggesting that the variability in clone size is related to its position. Cluster size was generally heterogeneous across samples (Figs. 5H-H’’), indicating that not all hepatoblasts contribute equally to the adult organ (Fig. 5I).

**Fig. 5.**
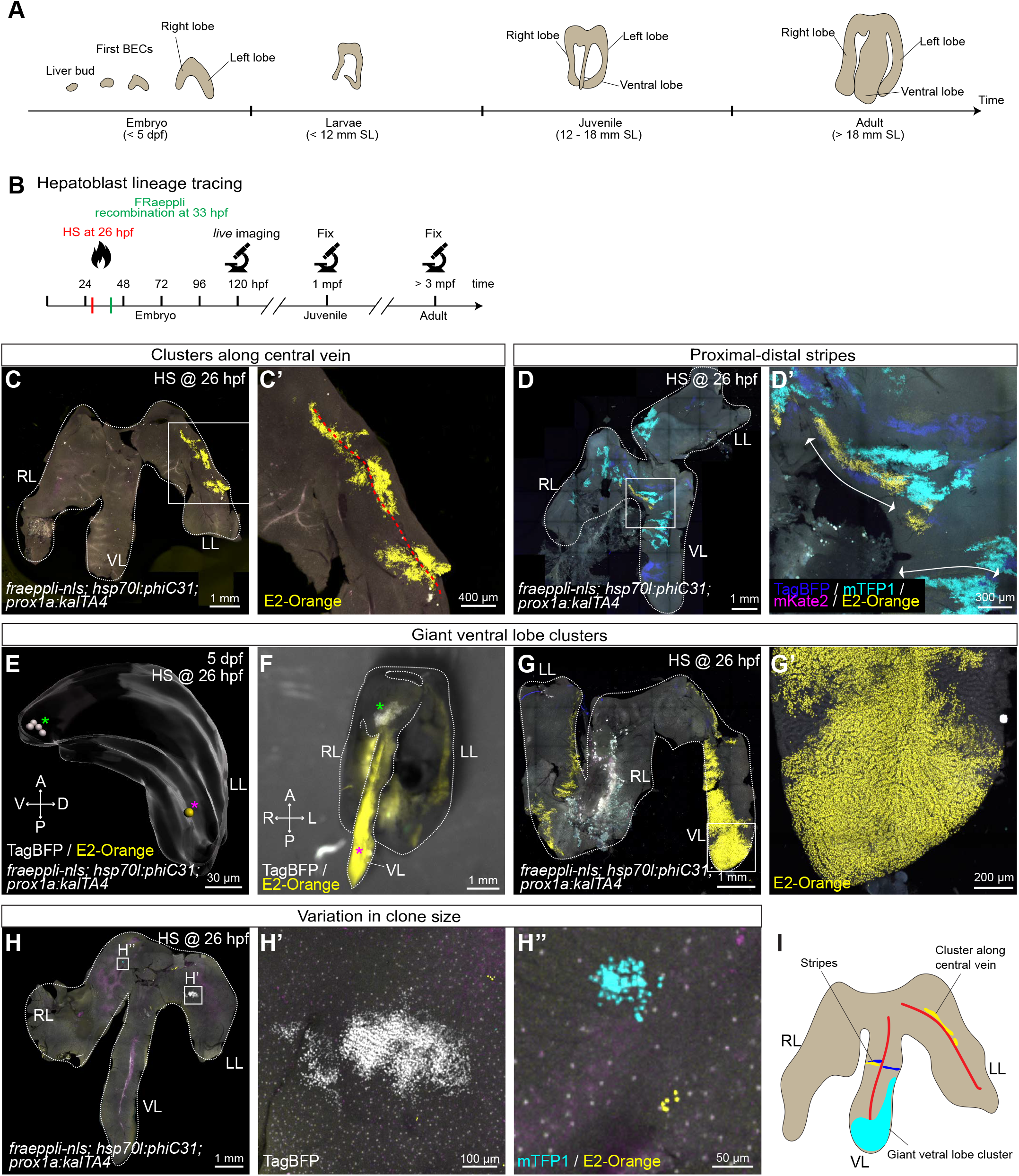
Lineage tracing reveals heterogeneous cluster topologies during postembryonic growth. (A) Schematic depicting key stages in postembryonic zebrafish liver development. (B) Experimental schematics of long-term lineage tracing experiments using *fraeppli-nls* embryos, inducing recombination by heat shock at 26 hpf to label hepatoblasts. At 120 hpf, embryos were screened by *live imaging* at the confocal microscope and only sparsely labelled embryos were raised and fixed in either juvenile or adult stages. (C-H) Recombined livers showed different cluster topologies: clusters along central veins (C-C’) (N=9, n=9), proximal-distal stripes (D) (N=9, n=23) or giant clusters in the ventral lobe in adult (F-G’) (N=9, n=3). Large clusters in the ventral lobe can originate from one single-labelled cell at 5 dpf (E). (F) Stereomicroscope image showing the spatial location of the giant clone originating from a single recombined cell (H). Recombined livers show a range of cluster sizes from small (H’) to medium (H’’). (I) Schematics of characteristic cluster topologies in recombined livers. Red lines indicate the blood vessel orientation in the liver. A = anterior, P = posterior, R = right, L = left, RL = right lobe, LL = left lobe, VL = ventral lobe.

To distinguish whether cluster shape and size are inherent to progenitors or influenced by other, external factors, we performed a second set of experiments in which recombination was induced in differentiated hepatocytes. For this *fraeppli-nls; fabp10a:kalTA4* were crossed to *hsp70l:phiC31* and administered a heat shock at 4 dpf for sparse labelling and 42 juvenile and 31 adult livers were analysed. In 16% of adult livers clusters extended along the large central vein (Fig. S4D-D’) and 32% exhibited proximodistal stripes (Fig. S4E-E’), consistent with the results of recombined hepatoblasts. At the same time, we could not detect any giant cluster in the ventral lobe. However, within the juvenile samples induced at 4 dpf, we identified in a single instance, representing 2.3%, a giant cluster in the ventral lobe, comprising 1.7% of the total liver volume (Fig. S4F-F’). Hence, suggesting that hepatocytes can also contribute giant clusters to the ventral lobe. Due to the lower sample number, compared to the hepatoblast tracing, we did not observe this topological class in adults induced at 4 dpf. Interestingly, we did not observe any proximodistally-striped clusters in juvenile livers, suggesting that growth along this axis predominantly occurs in late juvenile stages.

To assess the specificity of *hsp70l:phiC31* expression in non-induced embryos, we examined 100 juvenile or adult control livers and concluded that unspecific recombination is rare (Fig. S4 H-J). Overall, these findings suggest that cluster shapes and growth patterns are independent of the intrinsic proliferation potential of single hepatoblasts, or the differentiation state of the recombined cell, and instead are the result of extrinsic growth signals.

To investigate hepatic growth dynamics, we examined whether proliferating cells are evenly distributed or spatially enriched during development. Interestingly, from 84 hpf onwards proliferating cells are located significantly closer to the surface than the centre of the liver (Fig. S5A-B), suggesting that growth is enhanced at the organ periphery. Measuring the distance to the nearest neighbours between proliferating hepatocytes and BECs or across each cell population, revealed that average distances between proliferating cells are greater than between all cells of the given population (Fig S5C-D), rejecting the possibility of proliferation clusters.

### Polyploid cells appear transiently during postembryonic liver growth

Postembryonic growth (Fig. S6A) is poorly understood across organs and species. Monitoring liver growth by comparing liver to body weight (Fig. S6B-C) between late larval to adult stages revealed a two-phase process: the liver grows faster than the body in juveniles until it attains a stable liver to body weight ratio once the fish reach adulthood, with an average of 5.5% for female and 3.5% for male zebrafish (Fig. S6D). However, the proportion of liver to body weight in juveniles is higher, on average 8.6%, and more variable (2.1 -22.2%; Fig. S6D), likely reflecting substantial organ growth including the formation of an additional lobe.

From a mechanistic point of view, rapid growth could be achieved by increasing cell size, cell ploidy and proliferation, including *de novo* tissue extensions. First addressing whether polyploid hepatocytes are present in the zebrafish liver, particularly during periods of rapid organ growth, we analysed whole-mount livers of *fabp10a:GFP; tp1:H2B-mCherry* counterstained with 4’,6-diamidino-2-phenylindole (DAPI). At the larval stage, at 14 dpf, shortly after liver function commences, the volume of hepatocyte nuclei was variable, and rare binucleated hepatocytes appeared in 50% of embryos (Figs. 6A-A’’). Subsequently, in juveniles, we observed two scenarios: multinucleated hepatocytes and enlarged nuclei (Figs. 6B-B’), with an up to 10-fold volume increase and elevated DAPI intensity compared to average hepatocyte nuclear values (Figs. 6C-D), indicating that zebrafish hepatocytes enter a polyploid state. Contrary to larval livers, all juvenile samples exhibit enlarged nuclei and multinucleated cells at high frequency in all three lobes. Next, we assessed whether polyploid hepatocytes are maintained, become more abundant, or disappear with age. Surprisingly, we detected only a single group of three polyploid nuclei in one of five adult livers (Figs. 6E-G), showing that hepatocyte polyploidy is transient, as it peaks with the massive growth phase of the zebrafish liver and declines with maturation (Fig. 6H).

**Fig. 6.**
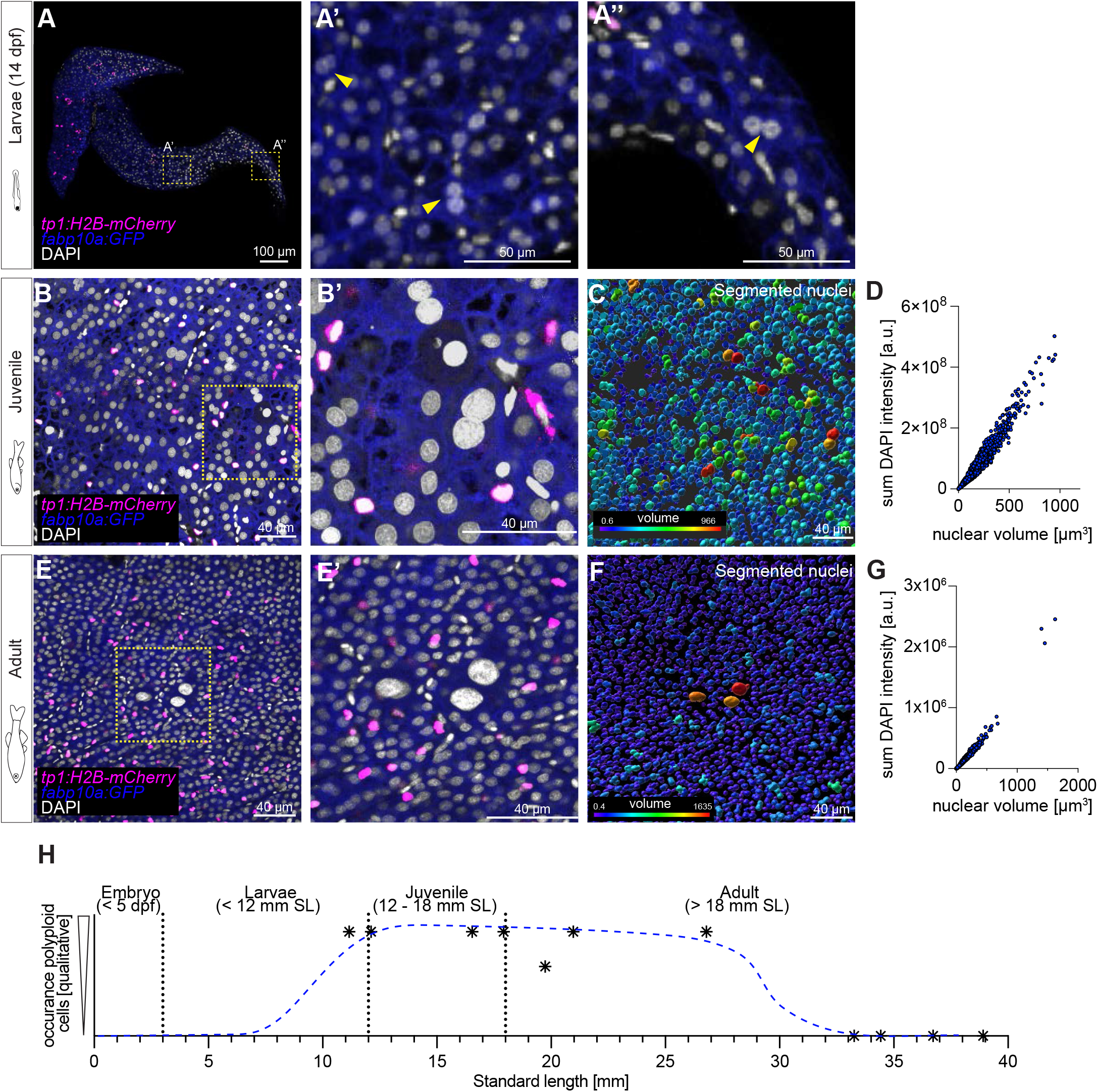
Polyploid cells appear transiently in hepatic postembryonic growth in zebrafish. (A) Whole-mount of a 14 dpf zebrafish liver, displaying sparse multinucleated hepatocytes (N=1, n=2; yellow arrowheads indicate binucleated cells). (B) 5 µm projection of a region of a juvenile liver. Fish SL = 11,16mm (N=3, n=6). (C) Segmentation shows variable nuclear volumes, which correlate with the sum intensity of DAPI, indicating that bigger nuclei have a higher amount of DNA (D). (E) 5 µm projection of an adult liver region (N=3, n=3). (F) Segmented nuclei show only sparse variability in volume, with few bigger nuclei. Nuclear volume correlates with sum intensity of DAPI (G). (H) Schematics representing the transient appearance of polyploid cells over time; blue trajectory is manually approximated based on qualitative analysis.

### Liver morphology and clonal growth patterns reveal distinct ventral lobe formation

Assessing whether other mechanisms besides ploidy may contribute to postembryonic liver growth, such as extension by additional organ parts, we turned our attention to the *de novo-* forming ventral lobe, because of the unique giant clusters and their distinct clonal growth behaviour. The ventral lobe is described to arise from the ventral part of the liver [21], yet experimental data concerning its origin and formation are missing. Therefore, to relate the origin of the giant clusters to ventral lobe formation, we carefully examined liver morphology throughout postembryonic growth in the 62 larval and juvenile livers collected for the lineage tracing studies. Based on morphological characteristics, ventral lobe formation across postembryonic growth was divided in six phases (Fig. 7A). During stage I, the liver consists mainly of the right and left lobe, with the latter exhibiting a slight bulge towards the ventral midline (Fig. 7B). During stage II, the ventral lobe has started to form and identifies with a very thin structure which originates in the more posterior half of the left lobe (Fig. 7C). The position of ventral lobe outgrowth shifts to the more anterior part of the left lobe during stage III, while it still maintains its long thin and flat appearance (Fig. 7D). The tip of the ventral lobe starts to round up and expands in a radial manner during stage IV and at the same time the base of the ventral lobe broadens, strengthening its connection at the ventral most part of the liver (Fig. 7E). Thereafter during stage V, the ventral lobe increases in size expanding laterally (Fig.7F), reaching its final width in late juvenile stages or early adulthood (stage VI, Fig.7G).

**Fig. 7.**
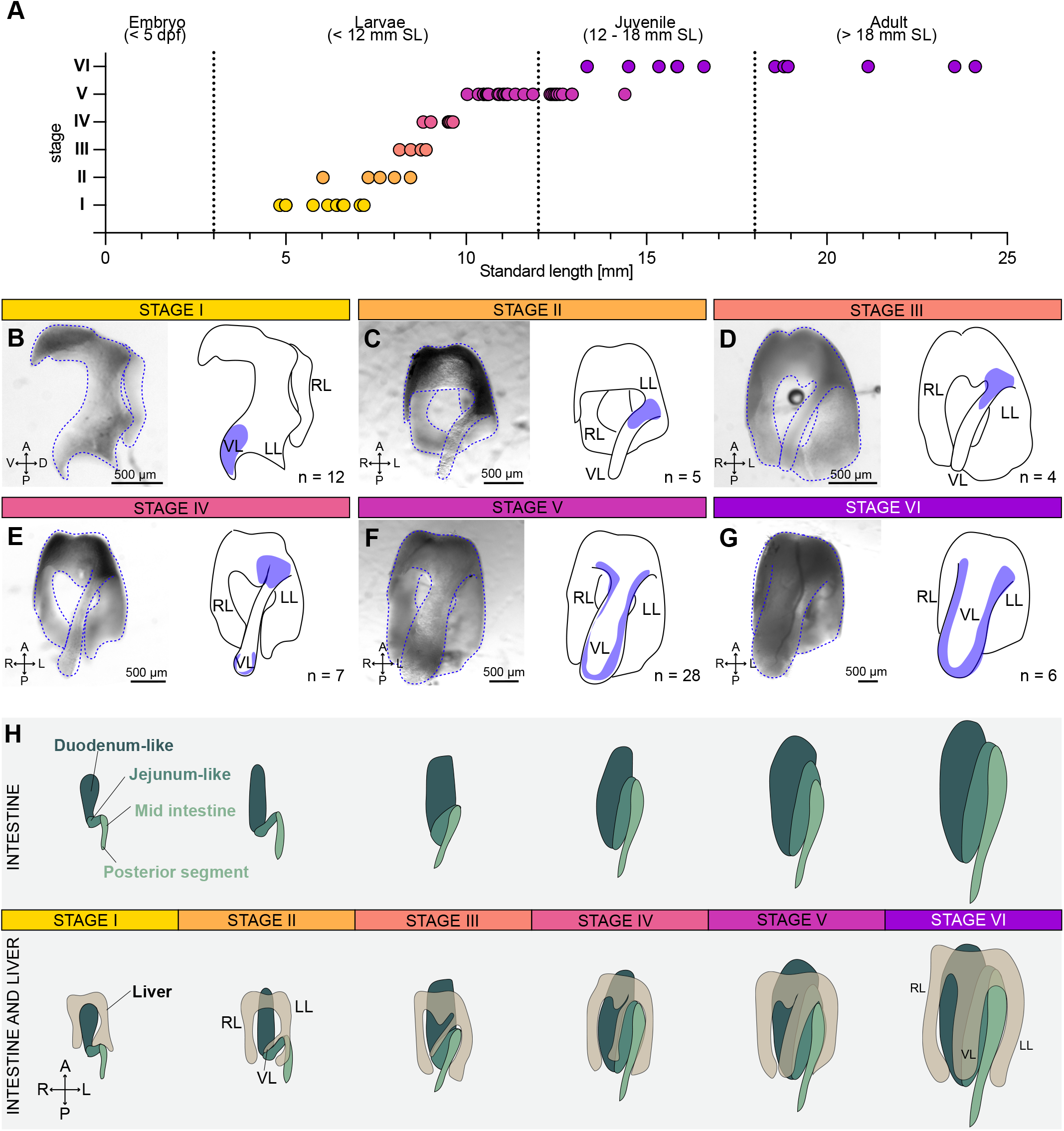
Ventral liver lobe formation during postembryonic growth. (A) The six steps of ventral liver lobe formation correlate with fish standard length (SL). (B) Stage I: a small tissue extension at the tip of the left lobe is visible (n=12). (C) Stage II: a thin ventral lobe originates in the lower half of the left lobe (n=6). (D) Stage III: the thin ventral lobe shifts position towards the more anterior part of the left lobe (n=4). (E) Stage IV: the tip of the ventral lobe starts to expand (n=7). (F) Stage V: lateral-oriented expansion of the ventral lobe (n=28). (G) Stage VI: enlargement of all lobes in width (n=6). The blue areas in the schematics mark the region characteristic for the respective stage. (H) Schematic depicting the morphology of the liver in relation to the folding of the intestine in stages I-VI. A = anterior, P = posterior, R = right, L = left, RL = right lobe, LL = left lobe, VL = ventral lobe.

Using these morphological data as a reference, we turned to the long-term lineage tracing data asking whether the future ventral lobe would arise from the embryonic left lobe. We correlated characteristic ventral lobe cluster patterns in juvenile and adult livers, with corresponding clonal positions within the same fish acquired at 5 dpf. For instance, a 3-cell mKate2^+^ clone located at the periphery of the left lobe at 5 dpf (Fig. S7A), grew into a 1420-cell mKate2^+^ clone at the juvenile stage, exclusively localised in the ventral lobe (Fig. S7B). Furthermore, 36% of juvenile livers contained clusters displaying a similar orientation from the left lobe into the tip of the ventral lobe across stages (Fig. S7C,D). These distinct and stereotypic clone patterns strongly support our hypothesis that the ventral lobe grows out from the left lobe.

Interestingly, the most dramatic morphological changes occur during the larval to juvenile transition (stages I-IV), mirroring zebrafish metamorphosis, during which many organs transform and adopt adult characteristics, such as the gut [23]. Considering the overall morphological changes occurring during zebrafish metamorphosis, the morphogenesis of the intestine, in particular the appearance of the two intestinal bends, coincides temporally and spatially with the repositioning of the ventral lobe (Fig. 7H, S7E-P). The intestine and liver are in direct contact during those stages, with the ventral lobe situated directly on top of the first intestinal fold. Notably, when the gut starts folding, ventral lobe formation is initiated (stage I) (Fig. S7E-F). With progressive bending of the gut, the position of the outgrowing ventral lobe shifts anteriorly (stage II - V) (Fig. S7G-N). In parallel, as the ventral lobe expands laterally (stage VI), it is located directly above the intestinal fold (Fig. S7O-P). Finally, in adults, the three liver lobes almost entirely enwrap the intestine. These data suggest that gut and liver morphogenesis are coupled during metamorphosis (Fig. 7H), in line with the idea of stimulating cues from the intestine promoting the selective expansion of the ventral lobe, as mirrored by the spatial expansion dynamics of the giant clusters.

## Discussion

The contribution of multipotent progenitor cells to organ formation and the establishment of correct cell proportions is vital for building a functional organ. This study shows how heterogeneous hepatoblasts contribute to fate decisions and postembryonic liver growth (Fig. 8A-B), including de novo structures, enabling the formation of a functional liver architecture. We show that a precise 1:9 BEC to hepatocyte ratio within the liver is established and remains nearly constant into adulthood. Cell counts from single cell transcriptome studies of adult zebrafish match this ratio [40], corroborating that specific and constant cell proportions are pivotal for a functional architecture across all stages. Notably, this distinct ratio is already established before hepatic cells take up their function, allowing time for the morphogenetic differentiation of the intricate liver architecture, reminiscent of stem cell-based tissue homeostasis in the intestine where an early lineage commitment leading to a precise cell type ratio is thought to facilitate maturation prior to function [41].

**Fig. 8.**
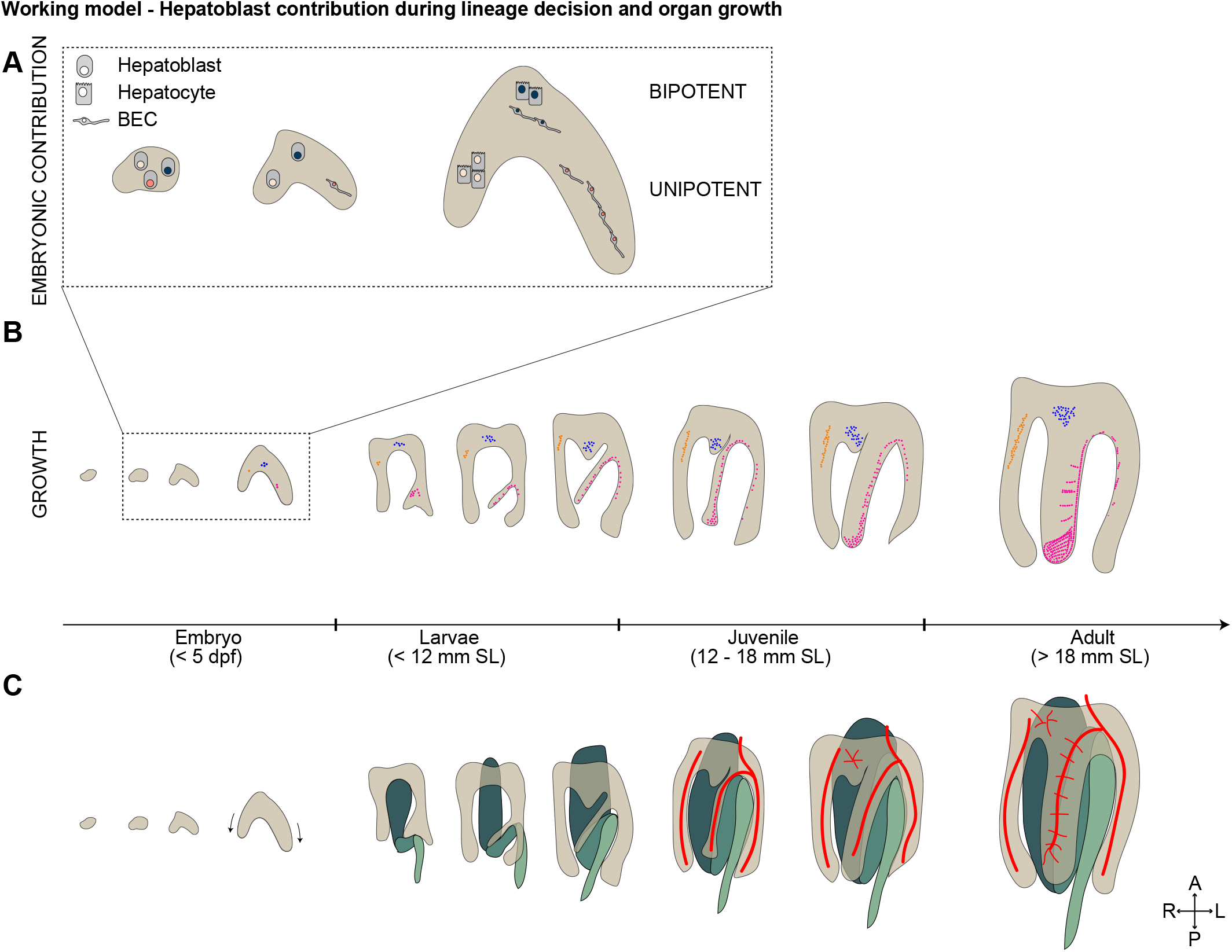
Working model of hepatoblast contribution to lineage decision and post embryonic growth. Schematics showing the current working models: (A) Uni- and bipotent hepatoblast contributions to hepatocytes and BECs following heterogeneous lineage decisions. (B) Hepatoblasts contribute with heterogeneous proliferation behaviours to postembryonic liver growth. Cells from the embryonic left lobe contribute to the ventral lobe, including the formation of giant clusters (magenta). (C) The liver morphology changes dramatically simultaneous to the intestinal bending occurring during postembryonic growth (green).

The establishment of a precise cell type ratio may be controlled by spatial signalling, such as Notch [42], given the seemingly even spacing of the first appearing BECs (Fig. 2F). Both in zebrafish and mice, Notch signalling plays a central role in BEC fate [16–18]. Similar to zebrafish, hepatocytes not only vastly outnumber BECs in homeostatic human livers but also in mice and rats [12,43]. With approximately 1:32 BECs to hepatocytes [12,44], these magnified proportions in the mouse may be explained by its tissue organisation in which not every hepatocyte directly connects to bile ductules like in zebrafish, so fewer BECs may be required for the overall network.

We combined *in silico* modelling with *in vivo* lineage tracing experiments to clearly show that lineage contributions of progenitor cells, and not differences in their proliferation rates, determine the given cell type ratio. These results unequivocally demonstrate the presence of bi- and unipotent hepatoblasts *in vivo* (Fig. 8A). Based on *in vitro* studies and lineage tracing of early foregut endoderm cells induced at E7.75, it has long been assumed that all hepatoblasts are bipotent and contribute to both cell lineages [11,13]. However, murine lineage tracing experiments of Lgr5^+^ hepatoblasts revealed functional heterogeneity of hepatoblasts [12], contributing equally to pure hepatocyte - or mixed hepatocyte and BEC clones adjacent to the portal triads [12]. Our analysis of clones throughout the liver using the pan-hepatoblast driver

*prox1a:kalTA4* revealed a larger proportion, 88.8%, of unipotent hepatocyte clones. Remarkably, we also uncovered a pure BEC clone indicating a rare population of unipotent BEC hepatoblasts which based on its large size, may have arisen from early BEC differentiation. This is supported by early cholangiocyte committed hepatoblasts, so far only been deduced computationally from single-cell RNA sequencing of murine hepatoblasts at E10.5 [12]. Our study shows uni- and bipotent hepatoblasts in zebrafish, demonstrating that heterogeneous lineage potential at the start of liver formation is conserved across species. We propose progenitor heterogeneity as a general strategy to set up lineage proportions in the liver, yet the underlying molecular mechanisms defining instructive signalling hierarchies [12] may differ, since an Lgr5 homologue seems to be missing in most teleost genomes, including zebrafish [45].

Growth dynamics of embryonic and especially postembryonic liver growth are poorly understood across species, including zebrafish. Here we show a steady decrease in BEC and hepatocyte proliferation, in line with the notion that high proliferation in undifferentiated tissues opposes lower proliferation rates in differentiated tissues in other organs [46]. Proliferating cells were enriched at the liver periphery, suggesting that external signals fuel tissue growth at the edge once differentiation ends and tissue architecture is established. Peripheral liver growth is regulated by β-catenin signalling in chicken [31], and Wilms tumor 1 from surrounding mesothelial cells promotes murine liver growth and lobe formation [29,30]. We show that the liver grows exponentially during metamorphosis and provide, to our knowledge the first evidence for polypoid hepatocytes in the zebrafish, well described in most mammals [47]. Both large nuclei and multinucleated cells are prominent in juvenile livers, yet rare in adults, which is markedly more similar to the homeostatic 20-40% polyploidy in humans compared to the 75-94% in mice [47,48]. Polyploidy represents an attractive developmental strategy to quickly increase organ size while maintaining or even elevating metabolic function due to higher DNA content and enlarged cell volume [49,50]. Nevertheless, the overall link between polyploidy and growth remains unclear, since inhibiting the formation of polyploid hepatocytes does not impair liver growth [51] and their relevance for liver regeneration remains controversial [52–54].

We provide the first qualitative study of postembryonic liver growth by tracing hepatoblast- and hepatocyte-progeny thereby identifying distinct growth patterns (Fig.8B). Remarkably, two cluster categories arranged along blood vessels, one paralleling the central vein and the other the perpendicularly organised sinusoids. This is in line with vessel-derived mitogens that may control hepatocyte proliferation, such as Wnt2, Angiopoetin and Hepatic growth factor, as well as signalling induced by mechanical stretching of endothelial cells by the incoming blood [55–58]. These cellular relationships further agree with the finding that hepatocytes and their newly arising daughter cells align parallel to sinusoids after hepatocyte damage [59] as well as cohesive and oriented growth at the organ surface during lobe development [60]. Lastly, the finding of giant clusters associated with ventral lobe growth is surprising, as it suggests growth dynamics and signals distinct from the two dorsal lobes. Strikingly, the correlation between adult clonal growth patterns and their embryonic origin demonstrates that the ventral lobe arises surprisingly from the embryonic left lobe and not the most ventral part, as previously suggested [21]. The overproportionate contribution of these giant clusters to the adult organ, therefore, likely reflects that only a few embryonic cells compared to the two dorsal lobes produce a lobe of similar size and is reminiscent of the clonal dominance observed in other tissues [27,61,62]. Given the stereotypic pattern of the giant clusters within the ventral lobe strongly indicates localised growth driving postembryonic organ morphogenesis. Signals could arise from the spatially and temporally correlating remodelling of the intestine and markedly the formation of its two bends (Fig. 8C). The intestine may provide physical constraints, metabolites and/or mitogens directing the characteristic growth of the ventral liver lobe. Inter-organ communication, such as for the *Drosophila* testes and intestine [63] represents an attractive strategy for guiding the distinct formation of the ventral liver lobe. Understanding differential modes of lobe formation is also highly relevant for regeneration studies in adults, since due to its accessibility, the ventral lobe is targeted for partial hepatectomy in zebrafish [64,65]. Yet, depending on the extent of injury the liver responds either with epimorphic or compensatory regeneration [32]. The underlying mechanisms remain elusive, thus further studies of postembryonic liver growth and in particular ventral lobe formation are pivotal.

In summary, we show that the heterogeneous lineage contribution of hepatoblasts is the predominant factor establishing the distinct cell proportions of the functional liver, while heterogeneous proliferation dynamics of individual progenitors establish organ size. We propose that both lineage and proliferation heterogeneity is not intrinsic hepatoblast property, but stochastic and influenced by signals from the microenvironment. Identifying the molecular nature of these signals, as well as the morphogenetic principles directing tissue architecture will aid in developing strategies promoting the endogenous capacity of the liver to restore a functional tissue organisation and instruct engineering of hepatic tissues *in vitro*.

## Acknowledgements

We thank the Ober group for discussion and comments on the manuscript. We are grateful Dr. F. Lemaigre for feedback on the manuscript and Dr. T. Piotrowski for invaluable support.. We thank the department of experimental medicine (AEM) in Copenhagen for expert fish care. We gratefully acknowledge the DanStem Imaging Platform (University of Copenhagen) for support and assistance in this work.

## Data availability

The authors declare that all data supporting the findings of this study are available within the article and its supplementary information files or from the corresponding author upon reasonable request. The image data generated and analysed in this study are available from the corresponding author upon reasonable request.

## Material and Methods

### Zebrafish husbandry

Zebrafish *(Danio rerio)* embryos and adults were kept according to standard laboratory conditions [66]. All experiments were performed according to ethical guidelines approved by the Danish Animal Experiments Inspectorate (Dyreforsøgstilsynet).

The following transgenic strains were used: *tgBAC(prox1a:kalTA4)*^*uq3bh*^ [67], *tg(kdrl:EGFP)*^*s843*^ [68], *tg(EPV*.*TP1-Mmu*.*Hbb:hist2h2l-mCherry)*^*s939*^ [69], *tg(−2*.*8fabp10a:EGFP)*^*as3*^ [70], *tg(tp1-MmHbb:EGFP)*^*um14*^ [71], *tg(fraeppli-nls)*^*cph1-3, cph9*^ [34],*tg(fabp10a:kalTA4; cryaa:Venus)*^*cph8*^ [34],*tg(hsp70l:phiC31-integrase, he1a:lyn-Citrine)*^*cph7*^ [34].

### FRaeppli activation

In all experiments, FRaeppli recombination was induced conditionally using *tg(hsp70l:phiC31-integrase, he1a:lyn-Citrine)*^*cph7*^ [34].To induce *phiC31* expression, embryos were subjected to a 30 min heat shock at 39°C. Control embryos were kept at 28°C. To avoid undesired recombination due to temperature fluctuations, embryos were stored at strict temperature control, including double Styrofoam boxes for transport of recombined embryos. To express FRaeppli colours in hepatoblasts and subsequently hepatocytes and BECs or only hepatocytes expression was controlled by either *tgBAC(prox1a:kalTA4)*^*uq3bh*^ or *tg(fabp10a:kalTA4; cryaa:Venus)*^*cph8*^ respectively. We mainly used *fraeppli-nls*^*cph09*^, which recombines more sparsely than *fraeppli-nls*^*cph01-03*^.

### Imaging

All embryos were raised in E3 medium (5 mM NaCl, 0.17 mM KCl, 0.33 mM CaCl2, and 0.33 mM MgSO4). Medium was supplemented with 0.2 mM 1-phenyl 2-thiourea (PTU, Sigma-Aldrich) for embryos intended for *live* imaging. During *live* imaging embryos and larvae were immobilized in 0.8% low melting agarose and anesthetized with Tricaine (164 mg/L; MS-222, Sigma-Aldrich) dissolved in E3/PTU.

To identify sparsely recombined *fraeppli-nls* embryos at 120 hpf, 10-40 embryos were mounted and screened at the confocal microscope for the presence of recombined cells, using spectral imaging. Imaging time was kept as short as possible, and embryos were released from agarose immediately after imaging.

300 µm vibratome sections or whole juvenile and adult livers were cleared using SeeDB2G [72]. Livers of zebrafish larvae with a SL smaller than 8 mm were directly mounted in Omnipaque350 (Sigma-Aldrich, Histodenz) for imaging. Fixed *fraeppli-nls* embryos were embedded in VectaShield (VWR, VECTH-100) for imaging. Imaging was performed using LSM 780 and 880 confocal microscopes equipped with PMT detectors for sequential imaging and a spectral detector (GaAsP-PMT, 32 channels, 410-694 nm range, 8.9 nm bandwidth) for spectral acquisition. 5-colour sequential imaging of adult recombined livers was performed with a Leica Stellaris confocal microscope equipped with a tuneable white-light laser, 488 and 448 nm diodes and four PMT/HyD detectors. Adult recombined FRaeppli livers were imaged to 500 µm in depth using the 10x objective. Detailed parameters for sequential and spectral imaging of FRaeppli-NLS modes are described in Caviglia et al [34]. Stained embryos embedded in Benzyl Alcohol-Benzyl Benzoate (BABB) (ratio 1:2) were imaged with a Leica SP8 confocal microscope. Brightfield and fluorescent overview images of recombined *fraeppli-nls* livers were acquired at a Leica Stereomicroscope (Leica, M205 FCA) equipped with a CCD microscope camera (Leica, DFC7000 GT).

### Immunostaining

Embryos and larvae were fixed with 4% PFA at 4°C overnight. For adult liver staining, the liver was embedded in 4% agarose and cut into 200 µm sections using a vibratome. Tissue sections were stained in 9 well glass plates. Immunostainings were performed as previously described [73]. In short, for nuclear staining, embryos were permeabilized using DNase I (ThermoFisher Scientific) treatment for 45 min at 37°C and then blocked in 10% donkey serum (Jackson Immunoresearch, 017-000-121) with 1% PBS-Triton X-100. Primary antibodies were incubated with 1% PBS-Triton X-100, 1% Dimethylsulfoxide (DMSO) and 10% donkey serum at 4°C for one or two nights slowly shaking.

Primary antibodies included: α-Hnf4α (goat polyclonal antibody, 1:100, Santa Cruz, sc-6556); α-mCherry (rat monoclonal antibody, 1:1000, Invitrogen, m11217), α-Prox1 (rabbit polyclonal antibody, 1:500, Angiobio, 11-002P) and α-2F11 (mouse monoclonal antibody, 1:1000, gift from Julian Lewis). After washing in 0.1% PBS-Triton X-100, secondary antibodies were incubated with 1% PBS-Triton X-100, 1% DMSO and 10% donkey serum at 4°C for one to two nights.

Secondary antibodies included: donkey-anti-goat 488 (1:200, Jackson Immunoresearch, 705-545-147), donkey-anti-rat-cy3 (1:500, Jackson Immunoresearch, 715-166-150), goat-anti-rabbit 647 (1:500, Jackson Immunoresearch) and donkey-anti-mouse 647 (1:500, Jackson Immunoresearch, 715-605-151). For DNA staining, samples were incubated with DAPI (1:100-500, Sigma, D9542) or TO-PRO-3 (1:50000, ThermoFisher) for 1 hour at room temperature or overnight at 4°C. Samples were washed in 0.1% PBST and dehydrated in methanol.

### Processing juvenile and adult zebrafish

Juvenile and adult zebrafish were fixed as whole fish with 4% PFA at 4°C for more than 20 hours using a rotator. To allow PFA penetration into deep tissues, the skin of the abdominal wall was opened prior to fixation. After fixation, samples were washed in PBS for at least 1h at room temperature using a rotator. An overview picture was taken of each fixed fish to determine the SL [22] by measuring the distance from the mouth to the start of the fin with Fiji. Whole fish weight was assessed using a precision balance (Sartorius, Qunintx). The liver was dissected and weighed again in a pre-weighed Eppendorf tube filled with 1ml of PBS to avoid errors in the low weight range of the scale. For confocal imaging, livers were mounted flat between two coverslips or on a slide with two imaging spacers. We observed a large variation in liver shape with respect to lobe length and size, although the ventral lobe is consistently flatter than the two dorsal ones. In the majority of fish, the left and right lobe are merged dorsally at the posterior tips of the lobes and show connected vascular networks, whereas others display three independent lobes only connected at the most anterior part suggesting that liver growth and organ morphogenesis is plastic. When the tissue was connected between the tips of the left and right lobes, the connecting tissue was carefully cut with forceps to ensure flat mounting.

### Image analysis and cell segmentation

Images were processed with Bitplane Imaris and Zeiss ZEN software. As default, maximum intensity projections are displayed, unless indicated differently in the figure legends. Lambda scans were unmixed using HyperSpectral Phasor software (HySP) (version 0.9.10) [74]. A workflow describing the unmixing of FRaeppli FPs with HySP has been published in detail [34] For all embryonic images, the automated spot detection function in Imaris was used for quantification of cell numbers. EdU-positive cells of each cell type have been identified by segmenting EdU-positive nuclei and filtering for expression level of either Hnf4 (hepatocyte) or *tp1:mCherry* (BEC). The liver volume was segmented by manual surface generation in Imaris based on morphological difference of the liver tissues compared to surrounding tissues. Distances of cells to the liver surface was extracted in Imaris by using Spot-to-Surface distance. Distances of cells to nearest neighbours were also automatically obtained after segmentation in the latest version of Imaris, Imaris 9.9. To determine ploidy, nuclear volume was segmented using surface function in Imaris and Sum Intensity of DAPI measured within the segmentation.

To determine the ratio of endodermal cell types in adult tissues we investigated four juvenile and two adult livers stained with DAPI. Of each juvenile liver and one adult, we extracted at least three regions of interest (ROIs) of whole-mount images acquired with a 40x Objective with a size of 1000 × 1000 × 50 µm and segmented the nuclei. In addition, one adult liver was sectioned in 300 µm thick agarose sections using the vibratome and at least two to three ROIs from each of the three sections were selected. When available, ROIs taken from different lobes were included. For BEC nuclei we used the *tp1:H2B-mCherry* signal in the 564 nm channel for segmentation. For quantifying hepatocytes, DAPI signal was segmented, and filtered against high intensity cells to exclude blood cells and for expressing GFP from *fabp10a:GFP* transgene. Although we observed that the localization of the endogenous fabp10 signal varied between samples, from a homogeneous cytoplasmic signal to a more peripheral localization, the signal was sufficient to identify hepatocyte nuclei by determining the overlapping signal of DAPI with GFP.

For embryonic samples, *fraeppli-nls*-marked cells were segmented using the Imaris spot function. The xyz-coordinates were extracted and subsequently used for the mathematical analysis of clone determination. BEC nuclei were identified by filtering spots for the expression of GFP (*tp1:EGFP*). In embryos without *tp1:EGFP*, the elongated nuclear shape was used as a proxy for assigning BEC fate. Only very elongated nuclei were categorized as BEC. In case of doubt, the clone was excluded from the analysis.

For adult recombined *fraeppli-nls* livers, images were binned in Fiji by factor two in x and y to reduce image size. Subsequently, clonal volume was quantified in Imaris using the surface function.

### EdU incorporation assay and proliferation analysis

To label proliferating cells, embryos were incubated with 400 µM 5-ethynyl-2’-deoxyuridine (EdU) (Invitrogen)/DMSO in fish water for 1 hour at 28°C. Dependently on the stage, different DMSO concentrations were used: 48 – 60 hpf: 15% DMSO; 72 hpf: 10 % DMSO; 84 – 144 hpf: 5% DMSO. Upon incubation, embryos were immediately fixed in 4% PFA. EdU labelled cells were detected with the Click-it EdU Imaging Kit Alexa Fluor 647 (Invitrogen). Proliferating cell numbers were assessed using the Imaris software.

### Mathematical model of embryonic liver development

For simulating liver development, three distinct phases were included: proliferating hepatoblasts (phase I), differentiation into hepatocytes and BECs (phase II), followed by hepatocyte and BEC proliferation (phase III).

### The algorithm follows the steps listed below

First, we start with an initial number of cells in the hepatoblast state (n=100), with all cells proliferating at the same cell cycle length. Second, once hepatoblasts reach a specific number of cells (n=200), differentiation into hepatocytes and BECs is initiated. Cell differentiation is always linked to cell division of hepatoblasts. For the four different models, distinct probabilities for hepatoblast differentiation into hepatocytes and/or BEC were defined.

|  | Model 1 | Model 2 | Model 3 | Model 4 | Model 5 | Model 6 |
| --- | --- | --- | --- | --- | --- | --- |
| p_divide [P → H + C] | 1 | 1 | 0.5 | 0.2 | 0.2 | 0.2 |
| p_divide [P → C + C] | 0 | 0 | 0 | 0.05 | 0.05 | 0 |
| p_divide [P → H + H] | 0 | 0 | 0.5 | 0.75 | 0.75 | 0.8 |

Third, once cells acquire the hepatocyte or BEC fate, they continue to divide while maintaining the acquired fate. The cell type specific division rates are defined for each model as follows, whereas 10 (a.u.) corresponds to 24 hours.

|  | Model 1 | Model 2 | Model 3 | Model 4 | Model 5 | Model 6 |
| --- | --- | --- | --- | --- | --- | --- |
| division rate_P | 1/10 | 1/10 | 1/10 | 1/10 | 1/10 | 1/10 |
| division rate_C | 1/10 | 1/15 | 1/10 | 1/10 | 1/15 | 1/10 |
| division rate_H | 1/10 | 1/10 | 1/10 | 1/10 | 1/10 | 1/10 |

The simulation is terminated when 3000 cells are reached.

For non-synchronized, random cell divisions, we used a Gillespie algorithm to calculate the division time for each individual cell. The model is a 3D agent-based model where a cell divides by placing a new (daughter) cell close to itself. The positions of the cells are continuously being updated with the method previously developed by Nissen and colleagues [75].

The code was generated in Python. For each model, ten simulations were performed, and a script in Matlab R2022a was used for statistical analysis and generation of the plots. Python codes for the mathematical model are available upon request.

### Manual quantification of clones during embryonic lineage labelling

#### Determining clones

We assigned a rule to manually assign recombined cells of the same colours to clones. *Live* imaging of *fraeppli-nls; hsp70l:phiC31; prox1a:kalTA4* embryos showed that individual cells in the liver move a maximum of 20 µm apart and change position between 60 and 80 hpf (Fig. S2B), at a time when hepatic fate is already established. Considering the total volume of the liver, which increases 3.2-fold between 72 and 100 hpf, and nuclear movements of 20 µm maximum, we estimated that adjacent cells could move 70 µm apart between 60 and 80 hpf. We manually assigned clones to include all recombined cells of one colour within a radius of 70 µm distance. The spot measurement function was used in Imaris to determine intercellular distance. Cells defined as clones were grouped together. The position (xyz-coordinates) of each individual labelled cell was then subsequently extracted from Imaris for analysis of clonality. 12.4% of livers with very dense labelling at 100 hpf were excluded from the analysis due to difficulties in precise clone definition

#### Determining division frequency of clones

Division rates were calculated based on clone size. Clones with cell numbers outside exact divisions were rounded down, e.g., a 5-cell clone was categorized as 2 divisions. Rounding up did not change the overall division distribution.

### Quantitative clonal reconstruction and assessment of clonality

For a single population of progenitor cells undergoing stochastic fate choice (e.g. division and apoptosis), two key theoretical predictions are expected to hold.

Firstly, given a constant division rate k, the probability distribution for a cell population to divide n times within a specific time frame T, is predicted to follow a Poisson distribution: 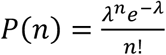, a distribution with λ*=kT* being the single adjustable parameter, equal to the average number of divisions in this time frame. This can therefore predict the full probability distribution for the number of cell divisions simply from the average properties (under the simplest assumption of a single population with a set division rate). Interestingly, we found that this simple assumption was enough to reproduce the global features of our experimental dataset (Fig. S3B).

Secondly, starting from a single labelled cell undergoing stochastic fate choices, the clone size distribution is expected to rapidly converge towards simple universal scaling laws [76]. In particular, for a 3D organ, the cumulative probability of clones consisting of m cells P(m) is predicted to adopt a simple exponential distribution at all time points: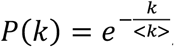, where <*k>*is the average clone size at this time point. This can be tested experimentally in datasets by plotting the clone size distribution in semi-log plot (Fig. S3A), in which an exponential distribution becomes a straight line. Interestingly, clone size distributions arising from manual reconstructions in the liver were shown to follow closely this trend.

To test this more systematically, we computationally reconstructed clones, while assessing from a statistical perspective the quality of the reconstruction. The core idea of this method [38] is to make use of the different colours in multispectral lineage tracing. In the simplest ideal scenario where all four colours are labelled with ¼ proportion each, we can then group clones according to specific rules (e.g. group all cells that are each with a certain distance d of each other in a colour-blind manner), and see whether we would have done “mistakes” under this criteria by grouping cells of different colours together in a clone. Said differently, if a specific method grouped cells of different colours together *x*%of the time, then the probability of grouping cells from two different clones of the same colour is 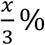.

We first assessed how variable were clonal inductions across different livers. Our dataset consisted of 97 fully reconstructed livers, each with the three-dimensional coordinates (x,y,z) of cells of each of the four colours. We first asked what the distribution of total number of labelled cells (of all colours) as per given liver and found that this was highly heterogeneous (Fig. S3C): half of the livers have less than 30 labelled cells, while 10% of livers have more than 100 cells (these long tails were markedly non-exponential and thus unlikely to occur by the stochasticity of cell division patterns, instead likely due to variable induction frequency).

We then computed for each labelled cell two metrics: the probability of finding at a distance r a cell of the same colour *P*_*same*_(*r*) and of a different colour *P*_*diff*_(*r*). In an ideal, sparse labelled scenario, cells should be all of the same colour in a certain critical radius *r*_*crit*_(where *P*_*same*_(*r*) is large and *P*_*diff*_(*r*) is small), while the probabilities should become identical at larger distance (as the colours of neighbouring clones are uncorrelated). We then calculated these distributions for different subsets of livers (the ones with less than 10 labelled cells, 20, 30 etc), with the expectation that livers with less labelled cells will show stronger clonality (Fig. S3D). Indeed, for livers with less than 10 labelled cells, we found that *P*_*same*_(*r*) and *P*_*diff*_(*r*)showed highly distinct peaks, with cells of a same colour being found less that 50μm apart, and of different colours above. Although these livers are likely to be biased towards small clones, this gives us an indication that around 50μm distance between cells might be a good criterion for clonal groupings. We further found that for livers with less than 50 cells, we could still find a clear distinction between the peaks of the two distributions, while this became less and less obvious when adding more densely-labelled livers (Fig. S3E). Interestingly, when grouping all cells of a given colour that were within 50μm of one another, and for livers with less than 40 cells, we found exponential clone size distributions close to the ones from the manual assessment of clonality, with average clone size of 4-5 cells.

### Statistics and reproducibility

Statistical analysis was performed using GraphPad Prism (version 9.3.1) or Matlab 2022a. The two-tailed Student’s t-test; n.s. p≥0.05 was used to calculate statistical significance. All data are presented as mean ± s.d. Unless indicated otherwise, n refers to sample size (e.g., individual embryos) and N refers to biological replicates.

## References

1. Arias MI, ALter HJ, Boyer JL, Cohen ED, Shafritz DA, Thorgeirsson SS, et al. The Liver: biology and pathology. 6th ed. Hoboken, NK: Wiley-Blackwell; 2020.

2. Lorent K, Yeo S-Y, Oda T, Chandrasekharappa S, Chitnis A, Matthews RP, et al. Inhibition of Jagged-mediated Notch signaling disrupts zebrafish biliary development and generates multiorgan defects compatible with an Alagille syndrome phenocopy. Development. 2004;131: 5753–5766. doi:10.1242/dev.01411.

3. Ota N, Shiojiri N. Comparative study on a novel lobule structure of the zebrafish liver and that of the mammalian liver. Cell Tissue Res. 2022;388: 287–299. doi:10.1007/s00441-022-03607-y.

4. Wang S, Miller SR, Ober EA, Sadler KC. Making It New Again: Insight Into Liver Development, Regeneration, and Disease From Zebrafish Research. Curr Top Dev Biol. 2017;124: 161–195. doi:10.1016/bs.ctdb.2016.11.012.

5. Poulain M, Ober EA. Interplay between Wnt2 and Wnt2bb controls multiple steps of early foregut-derived organ development. Development. 2011;138: 3557–3568. doi:10.1242/dev.055921.

6. Ober EA, Field HA, Stainier YR. From endoderm formation to liver and pancreas development in zebrafish. Mech Dev. 2003;120: 5–18. doi:10.1016/s0925-4773(02)00327-1.

7. Shin D, Monga SPS. Cellular and molecular basis of liver development. Compr Physiol. 2013;3: 799–815. doi:10.1002/cphy.c120022.

8. Cayuso J, Dzementsei A, Fischer JC, Karemore G, Caviglia S, Bartholdson J, et al. EphrinB1/EphB3b Coordinate Bidirectional Epithelial-Mesenchymal Interactions Controlling Liver Morphogenesis and Laterality. Dev Cell. 2016;39: 316–328. doi:10.1016/j.devcel.2016.10.009.

9. Germain L, Blouin MJ, Marceau N. Biliary epithelial and hepatocytic cell lineage relationships in embryonic rat liver as determined by the differential expression of cytokeratins, alphafetoprotein, albumin, and cell surface-exposed components. Cancer Res. 1988;48: 4909–4918.

10. Tanimizu N, Saito H, Mostov K, Miyajima A. Long-term culture of hepatic progenitors derived from mouse Dlk+ hepatoblasts. J Cell Sci. 2004;117: 6425–6434. doi:10.1242/jcs.01572.

11. Suzuki A, Zheng YW, Kondo R, Kusakabe M, Takada Y, Fukao K, et al. Flow-cytometric separation and enrichment of hepatic progenitor cells in the developing mouse liver. Hepatology. 2000;32: 1230–1239. doi:10.1053/jhep.2000.20349.

12. Prior N, Hindley CJ, Rost F, Meléndez E, Lau WWY, Göttgens B, et al. Lgr5+ stem and progenitor cells reside at the apex of a heterogeneous embryonic hepatoblast pool. Development. 2019;146: 1–14. doi:10.1242/dev.174557.

13. el Sebae GK, Malatos JM, Cone M-KE, Rhee S, Angelo JR, Mager J, et al. Single-cell murine genetic fate mapping reveals bipotential hepatoblasts and novel multi-organ endoderm progenitors. Development. 2018;145: dev168658. doi:10.1242/dev.168658.

14. Yang L, Wang WH, Qiu WL, Guo Z, Bi E, Xu CR. A single-cell transcriptomic analysis reveals precise pathways and regulatory mechanisms underlying hepatoblast differentiation. Hepatology. 2017;66: 1387–1401. doi:10.1002/hep.29353.

15. Gérard C, Tys J, Lemaigre FP. Gene regulatory networks in differentiation and direct reprogramming of hepatic cells. Semin Cell Dev Biol. 2017;66: 43–50. doi:10.1016/j.semcdb.2016.12.003.

16. Lorent K, Moore JC, Siekmann AF, Lawson N, Pack M. Reiterative use of the Notch signal during zebrafish intrahepatic biliary development. Developmental Dynamics. 2010;239: 855–864. doi:10.1002/dvdy.22220.

17. Zong Y, Panikkar A, Xu J, Antoniou A, Raynaud P, Lemaigre F, et al. Notch signaling controls liver development by regulating biliary differentiation. Development. 2009;136: 1727–1739. doi:10.1242/dev.029140.

18. Zhang D, Gates KP, Barske L, Wang G, Lancman JJ, Zeng XXI, et al. Endoderm Jagged induces liver and pancreas duct lineage in zebrafish. Nat Commun. 2017;8: 769. doi:10.1038/s41467-017-00666-6.

19. Ober EA, Lemaigre FP. Development of the liver: Insights into organ and tissue morphogenesis. J Hepatol. 2018;68: 1049–1062. doi:10.1016/j.jhep.2018.01.005.

20. Farber SA, Pack M, Ho S, Johnson ID, Wagner DS, Dosch R, et al. Genetic Analysis of Digestive Physiology Using Fluorescent Phospholipid Reporters. Science (1979). 2001;1385: 1385–1389. doi:10.1126/science.1060418.

21. Korzh S, Pan X, Garcia-Lecea M, Winata CL, Pan X, Wohland T, et al. Requirement of vasculogenesis and blood circulation in late stages of liver growth in zebrafish. BMC Dev Biol. 2008;8: 1–15. doi:10.1186/1471-213X-8-84.

22. Singleman C, Holtzman NG. Growth and Maturation in the Zebrafish, Danio Rerio : A Staging Tool for Teaching and Research. Developmental Dynamics. 2014;11: 396–406. doi:10.1089/zeb.2014.0976.

23. Parichy DM, Elizondo MR, Mills MG, Gordon TN, Engeszer RE. Normal table of postembryonic zebrafish development: Staging by externally visible anatomy of the living fish. Developmental Dynamics. 2009;238: 2975–3015. doi:10.1002/dvdy.22113.

24. Gao C, Zhu Z, Gao Y, Lo LJ, Chen J, Luo L, et al. Hepatocytes in a normal adult liver are derived solely from the embryonic hepatocytes. Journal of Genetics and Genomics. 2018;45: 173–175. doi:10.1016/j.jgg.2017.12.003.

25. Zhang W, Chen J, Ni R, Yang Q, Luo L, He J. Contributions of biliary epithelial cells to hepatocyte homeostasis and regeneration in zebrafish. iScience. 2021;24: 102142. doi:10.1016/j.isci.2021.102142.

26. Stanger BZ. Cellular homeostasis and repair in the mammalian liver. Annu Rev Physiol. 2015;77: 179–200. doi:10.1146/annurev-physiol-021113-170255.

27. Gupta V, Poss KD. Clonally dominant cardiomyocytes direct heart morphogenesis. Nature. 2012;484: 479–484. doi:10.1038/nature11045.

28. McKenna A, Findlay GM, Gagnon JA, Horwitz MS, Schier AF, Shendure J. Whole-organism lineage tracing by combinatorial and cumulative genome editing. Science (1979). 2016;353: aaf7907. doi:10.1126/science.aaf7907.

29. Ijpenberg A, Pérez-Pomares JM, Guadix JA, Carmona R, Portillo-Sánchez V, Macías D, et al. Wt1 and retinoic acid signaling are essential for stellate cell development and liver morphogenesis. Dev Biol. 2007;312: 157–170. doi:10.1016/j.ydbio.2007.09.014.

30. Onitsuka I, Tanaka M, Miyajima A. Characterization and Functional Analyses of Hepatic Mesothelial Cells in Mouse Liver Development. Gastroenterology. 2010;138: 1525–1535. doi:10.1053/j.gastro.2009.12.059.

31. Suksaweang S, Lin C-M, Jiang T-X, Hughes MW, Widelitz RB, Chuong C-M. Morphogenesis of chicken liver: identification of localized growth zones and the role of β-catenin/Wnt in size regulation. Dev Biol. 2003;266: 109–122. doi:10.1016/j.ydbio.2003.10.010.

32. Gao C, Peng J. All routes lead to Rome: multifaceted origin of hepatocytes during liver regeneration. Cell Regeneration. Springer; 2021. p. 2. doi:10.1186/s13619-020-00063-3.

33. Laudadio I, Manfroid I, Achouri Y, Schmidt D, Wilson MD, Cordi S, et al. A feedback loop between the liver-enriched transcription factor network and miR-122 controls hepatocyte differentiation. Gastroenterology. 2012;142: 119–129. doi:10.1053/j.gastro.2011.09.001.

34. Caviglia S, Unterweger IA, Gasiūnaitė A, Vanoosthuyse AE, Cutrale F, Trinh LA, et al. FRaeppli: a multispectral imaging toolbox for cell tracing and dense tissue analysis in zebrafish. Development. 2022;149. doi:10.1242/dev.199615.

35. Russell JO, Ko S, Monga SP, Shin D. Notch Inhibition Promotes Differentiation of Liver Progenitor Cells into Hepatocytes via sox9b Repression in Zebrafish. Stem Cells Int. 2019;2019: 1–11. doi:10.1155/2019/8451282.

36. Rulands S, Simons BD. Tracing cellular dynamics in tissue development, maintenance and disease. Curr Opin Cell Biol. 2016;43: 38–45. doi:10.1016/j.ceb.2016.07.001.

37. Dang Y, Rulands S. Making sense of fragmentation and merging in lineage tracing experiments. Frontiers in Cell and Developmental Biology. Frontiers Media S.A.; 2022. doi:10.3389/fcell.2022.1054476.

38. Sznurkowska MK, Hannezo E, Azzarelli R, Rulands S, Nestorowa S, Hindley CJ, et al. Defining Lineage Potential and Fate Behavior of Precursors during Pancreas Development. Dev Cell. 2018;46: 360–375. doi:10.1016/j.devcel.2018.06.028.

39. Koltowska K, Apitz H, Stamataki D, Hirst EMA, Verkade H, Salecker I, et al. Ssrp1a controls organogenesis by promoting cell cycle progression and RNA synthesis. Development (Cambridge). 2013;140: 1912–1918. doi:10.1242/dev.093583.

40. Morrison JK, DeRossi C, Alter IL, Nayar S, Giri M, Zhang C, et al. Single-cell transcriptomics reveals conserved cell identities and fibrogenic phenotypes in zebrafish and human liver. Hepatol Commun. 2022;6: 1–14. doi:10.1002/hep4.1930.

41. Tóth B, Ben-Moshe S, Gavish A, Barkai N, Itzkovitz S. Early commitment and robust differentiation in colonic crypts. Mol Syst Biol. 2017;13: 902. doi:10.15252/msb.20167283.

42. Sjöqvist M, Andersson ER. Do as I say, Not(ch) as I do: Lateral control of cell fate. Developmental Biology. Elsevier Inc.; 2019. pp. 58–70. doi:10.1016/j.ydbio.2017.09.032.

43. Blouin A, Bolender RP, Weibel ER. Distribution of organelles and membranes between hepatocytes and nonhepatocytes in the rat liver parenchyma. A stereological study. J Cell Biol. 1977;9: 441–455. doi:10.1083/jcb.72.2.441.

44. Willnow D, Benary U, Margineanu A, Vignola ML, Konrath F, Pongrac IM, et al. Quantitative lineage analysis identifies a hepato-pancreato-biliary progenitor niche. Nature. 2021;597: 87– 91. doi:10.1038/s41586-021-03844-1.

45. Aghaallaei N, Gruhl F, Schaefer CQ, Wernet T, Weinhardt V, Centanin L, et al. Identification, visualization and clonal analysis of intestinal stem cells in fish. Development (Cambridge). 2016;143: 3470–3480. doi:10.1242/dev.134098.

46. Ruijtenberg S, van den Heuvel S. Coordinating cell proliferation and differentiation: Antagonism between cell cycle regulators and cell type-specific gene expression. Cell Cycle. 2016;15: 196– 212. doi:10.1080/15384101.2015.1120925.

47. Celton-Morizur S, Desdouets C. Polyploidization of Liver Cells. Adv Exp Med Biol. 2010;646: 123– 135. doi:10.1007/978-1-4419-6199-0_8.

48. Wang M-J, Chen F, Lau JTY, Hu Y-P. Hepatocyte polyploidization and its association with pathophysiological processes Official journal of the Cell Death Differentiation Association. Cell Death Dis. 2017;8. doi:10.1038/cddis.2017.167.

49. Gentric G, Desdouets C, Celton-Morizur S. Hepatocytes Polyploidization and Cell Cycle Control in Liver Physiopathology. Int J Hepatol. 2012;2012: 1–8. doi:10.1155/2012/282430.

50. Kreutz C, MacNelly S, Follo M, Wäldin A, Binninger-Lacour P, Timmer J, et al. Hepatocyte ploidy is a diversity factor for liver homeostasis. Front Physiol. 2017;8. doi:10.3389/fphys.2017.00862.

51. Pandit SK, Westendorp B, Nantasanti S, van Liere E, Tooten PCJ, Cornelissen PWA, et al. E2F8 is essential for polyploidization in mammalian cells. Nat Cell Biol. 2012;14: 1181–1191. doi:10.1038/ncb2585.

52. González-Rosa JM, Sharpe M, Field D, Soonpaa MH, Field LJ, Burns CE, et al. Myocardial Polyploidization Creates a Barrier to Heart Regeneration in Zebrafish. Dev Cell. 2018;44: 433-446.e7. doi:10.1016/j.devcel.2018.01.021.

53. Matsumoto T, Wakefield L, Tarlow BD, Grompe M. In Vivo Lineage Tracing of Polyploid Hepatocytes Reveals Extensive Proliferation during Liver Regeneration. Cell Stem Cell. 2020;26: 34-47.e3. doi:10.1016/j.stem.2019.11.014.

54. Wilkinson PD, Delgado ER, Alencastro F, Leek MP, Roy N, Weirich MP, et al. The Polyploid State Restricts Hepatocyte Proliferation and Liver Regeneration in Mice. Hepatology. 2019;69. doi:0.1002/hep.30286.

55. Ding B-S, Nolan DJ, Butler JM, James D, Babazadeh AO, Rosenwaks Z, et al. Inductive angiocrine signals from sinusoidal endothelium are required for liver regeneration. Nature. 2010;468: 310– 315. doi:10.1038/nature09493.

56. Poisson J, Lemoinne S, Boulanger C, Durand F, Moreau R, Valla D, et al. Liver sinusoidal endothelial cells: Physiology and role in liver diseases. J Hepatol. 2017;66: 212–227. doi:10.1016/j.jhep.2016.07.009.

57. Lorenz L, Axnick J, Buschmann T, Henning C, Urner S, Fang S, et al. Mechanosensing by β1 integrin induces angiocrine signals for liver growth and survival. Nature. 2018;562: 128–132. doi:10.1038/s41586-018-0522-3.

58. Schoen JM, Wang HH, Minuk GY, Lautt WW. Shear stress-induced nitric oxide release triggers the liver regeneration cascade. Nitric Oxide. 2001;5: 453–464. doi:10.1006/niox.2001.0373.

59. Hoehme S, Brulport M, Bauer A, Bedawy E, Schormann W, Hermes M, et al. Prediction and validation of cell alignment along microvessels as order principle to restore tissue architecture in liver regeneration. PNAS. 2010;107: 10371–10376. doi:10.1073/pnas.0909374107.

60. Weiss MC, le Garrec J-F, Coqueran S, Strick-Marchand H, Buckingham M. Progressive developmental restriction, acquisition of left-right identity and cell growth behavior during lobe formation in mouse liver development. Development. 2016;143: 1149–1159. doi:10.1242/dev.132886.

61. Roan HY, Tseng TL, Chen CH. Whole-body clonal mapping identifies giant dominant clones in Zebrafish skin epidermis. Development. 2021;148: 1–11. doi:10.1242/DEV.199669.

62. Snippert HJ, van der Flier LG, Sato T, van Es JH, van den Born M, Kroon-Veenboer C, et al. Intestinal crypt homeostasis results from neutral competition between symmetrically dividing Lgr5 stem cells. Cell. 2010;143: 134–144. doi:10.1016/j.cell.2010.09.016.

63. Hudry B, de Goeij E, Mineo A, Gaspar P, Hadjieconomou D, Studd C, et al. Sex Differences in Intestinal Carbohydrate Metabolism Promote Food Intake and Sperm Maturation. Cell. 2019;178: 901-918.e16. doi:10.1016/j.cell.2019.07.029.

64. Oderberg IM, Goessling W. A protocol for partial hepatectomy in adult zebrafish. J Vis Exp. 2021;170. doi:10.3791/62349.

65. Sadler KC, Krahn KN, Gaur NA, Ukomadu C. Liver growth in the embryo and during liver regeneration in zebrafish requires the cell cycle regulator, uhrf1. PNAS. 2007;104: 1570–1575. doi:10.1073/pnas.0610774104.

66. Westerfield M. The zebrafish book. A guide for the laboratory use of zebrafish (Danio rerio). 4th ed. Univ. of Oregon Press, Eugene; 2000.

67. Koltowska K, Lagendijk AK, Pichol-Thievend C, Fischer JC, Francois M, Ober EA, et al. Vegfc Regulates Bipotential Precursor Division and Prox1 Expression to Promote Lymphatic Identity in Zebrafish. Cell Rep. 2015;13: 1828–1841. doi:10.1016/j.celrep.2015.10.055.

68. Jin SW, Beis D, Mitchell T, Chen JN, Stainier DYR. Cellular and molecular analyses of vascular tube and lumen formation in zebrafish. Development. 2005;132: 5199–5209. doi:10.1242/dev.02087.

69. Ninov N, Borius M, Stainier DYR. Different levels of Notch signaling regulate quiescence, renewal and differentiation in pancreatic endocrine progenitors. Development. 2012;139: 1557–1567. doi:10.1242/dev.076000.

70. Her GM, Chiang CC, Chen WY, Wu JL. In vivo studies of liver-type fatty acid binding protein (L-FABP) gene expression in liver of transgenic zebrafish (Danio rerio). FEBS Lett. 2003;538: 125– 133. doi:10.1016/S0014-5793(03)00157-1.

71. Parsons MJ, Pisharath H, Yusuff S, Moore JC, Siekmann AF, Lawson N, et al. Notch-responsive cells initiate the secondary transition in larval zebrafish pancreas. Mech Dev. 2009;126: 898– 912. doi:10.1016/j.mod.2009.07.002.

72. Ke MT, Nakai Y, Fujimoto S, Takayama R, Yoshida S, Kitajima TS, et al. Super-Resolution Mapping of Neuronal Circuitry With an Index-Optimized Clearing Agent. Cell Rep. 2016;14: 2718–2732. doi:10.1016/j.celrep.2016.02.057.

73. Thestrup MI, Caviglia S, Cayuso J, Heyne RLS, Ahmad R, Hofmeister W, et al. A morphogenetic EphB/EphrinB code controls hepatopancreatic duct formation. Nat Commun. 2019;10: 1–15. doi:10.1038/s41467-019-13149-7.

74. Cutrale F, Trivedi V, Trinh LA, Chiu CL, Choi JM, Artiga MS, et al. Hyperspectral phasor analysis enables multiplexed 5D in vivo imaging. Nat Methods. 2017;14: 149–152. doi:10.1038/nmeth.4134.

75. Nissen SB, Perera M, Gonzalez JM, Morgani SM, Jensen MH, Sneppen K, et al. Four simple rules that are sufficient to generate the mammalian blastocyst. PLoS Biol. 2017;15. doi:10.1371/journal.pbio.2000737.

76. Klein AM, Simons BD. Universal patterns of stem cell fate in cycling adult tissues. Development. 2011;138: 3103–3111. doi:10.1242/dev.060103.

