## Supplemental Figures for "Lineage tracing identifies heterogeneous hepatoblast contribution to cell lineages and postembryonic organ growth dynamics"

**Fig. S1. Distinct BEC and hepatocyte proportions: predictive *in silico* modelling of development and *in vivo* cell type quantification of postembryonic stages.** (A-C) Mathematical models simulating hepatoblast differentiation, based on heterogeneous hepatoblast potentials (A,B) or differential proliferation times (C; n=10). (D,F) 10 µm section of juvenile (D) and adult (F) livers stained for *fabp10a:GFP* (hepatocytes), *tp1:H2B-mCherry* (BECs) and DAPI (nuclei). (E) Relative distribution of BECs and hepatocytes in juvenile liver (n=4, n=4). (G) Relative distribution of BECs and hepatocytes at the organ centre or periphery in adult livers (N=1, n=1).

**Fig. S2. Defining parameters for lineage tracing experiments using the FRaeppli-NLS system.** (A) 10 µm projection of an adult liver section stained for Prox1 (magenta) and Anxa4 (green), white arrowheads indicate Prox1$^+$ BEC nuclei. The Prox1 signal was filtered using a median filter with a 3 pixel kernel for better visualization. (n=3 sections). (B) Schematic representation of the stepwise activation times of the *fraeppli* transgene. (C) Quantification of total liver cell numbers, encompassing hepatocytes and BECs, during development. (D-E) *fraeppli-nls* embryo showing only TagBFP and mTFP1 expression at 60 hpf (D), and expression of all four FRaeppli fluorescent proteins at 120 hpf (n=4). (F) Timelapse of TagBFP$^+$ and mTFP1$^+$ cells using spectral imaging of the liver upon heat shock induction at 9 hpf (N=2, n=3). Some neighbouring cells stay close together (magenta arrow), while others move up to 20 µm apart (green arrows). (G) Assignment criteria for manual clone definition.

**Fig. S3. Quantitative assessment of clonality.**
(A) Clone size distribution for different colours represented in a semi-log plot, shows highly consistent values and good fit to a simple exponential distribution (black line), expected for a single population undergoing stochastic division. (B) Number of cell divisions of manually-defined clones fit a Poisson distribution (black line) as expected for stochastic divisions (C) Cumulative probability of a certain number of labelled cells per liver (N=6, n = 97), showing that most livers have less than 50 labelled cells, but with heavy tails (10%-20%) of highly induced livers. (D) Probability that a given cell had a neighbouring cell with the same (red line) or a different colour (black line). Plots show subsets of the data that included livers with a total number of less than 10, 40, 50 or 100 labelled cells. Both distributions show high overlap for highly induced livers, which signifies poor clonality. For distances of less than 50 μm and livers with less than 40 cells, the ratio of "same" to "different" colour is high, meaning that nearby

cells of the same colour are unlikely to be non-clonal. (E) Manually determined size distribution of clones (black line) plotted together with different reconstructed clone size distributions. Different lines correspond to the regrouping of neighbouring cells of the same colour in the same clone if present within defined radii. Plots show subsets of the data that included livers with a total number of less than 10, 40, 50 or 100 labelled cells.

**Fig S4. Lineage tracing reveals a heterogeneous contribution of hepatocytes during postembryonic growth.** (A) Adult liver displaying clones along the central vein; recombination was induced in hepatoblasts at 26 hpf (N=4, n=10). (B-C) Adult livers exhibiting giant clusters in the ventral lobe (N=9, n=3). (D) Adult liver with a cluster along a central vein (N=4, n-5) and (E) clusters oriented in lateral stripes (N=4, n=10) upon recombination induced in hepatocytes. (F) Giant clusters in the ventral lobe are also apparent in juvenile livers when labelling was induced at 4 dpf in hepatocytes (N=5, n=1). (G) Confocal section showing the *kdrl:GFP*$^+$ sinusoidal architecture in the adult liver counterstained with DAPI (N=1, n=2). (H-J) 84% non-induced control livers of long-term lineage tracing experiments showed no recombined cells (H; N=15, n=84) and the majority of recombined samples (N=15, n=100) show only one recombined clone of a few labelled cells (I-J; N=15, n=16).

**Fig S5. Peripheral growth during liver development.** (A-B) Distribution of nuclear distance to the liver surface displayed for hepatocytes and EdU$^+$ hepatocyte (A), and BECs and EdU$^+$ BECs (B) (N=2, n≥6). (C-D) Distribution of nuclear distance to the nearest neighbour (NN) shown for hepatocytes and EdU$^+$ hepatocytes (C) and BECs and EdU$^+$ BECs (D) (N=2, n≥8).

**Fig. S6. Hepatic growth dynamics of postembryonic zebrafish.** (A) Fish standard length (SL) plotted against fish age. (B-C) Fish weight (B) and liver weight (C) increases with SL represented in a semi-log plot. (D) Liver to body weight ratio during postembryonic growth is constant in adult fish. (N>10, n≥300). Gender of the corresponding samples is colour-coded: male (blue), female (pink) and ND (green).

**Fig S7. Postembryonic ventral lobe formation.** (A-B) Confocal images of the same liver showing the embryonic left liver lobe at 5 dpf with a 3-cell mKate2$^+$ clone (A) and at juvenile stage (SL = 14.4 mm) including a continuous Kate2+ clone in the ventral lobe (N=1, n=1). (C-D) Juvenile livers (C – SL=8.46 mm and D – SL=10.93 mm) with connected clusters that are oriented along the tissue edge and spread through the left and the ventral lobe. (E-P) Brightfield images of stage I-VI livers in loco within the fish (E,G,I,K,M,O) or dissected out (F,H,J,L,N,P). In (M) the liver is removed and the gut bend is visible. Arrows indicate cluster growth direction. (N=4, n=14). A = anterior, P = posterior, R = right, L = left, RL = right lobe, LL = left lobe, VL = ventral lobe.

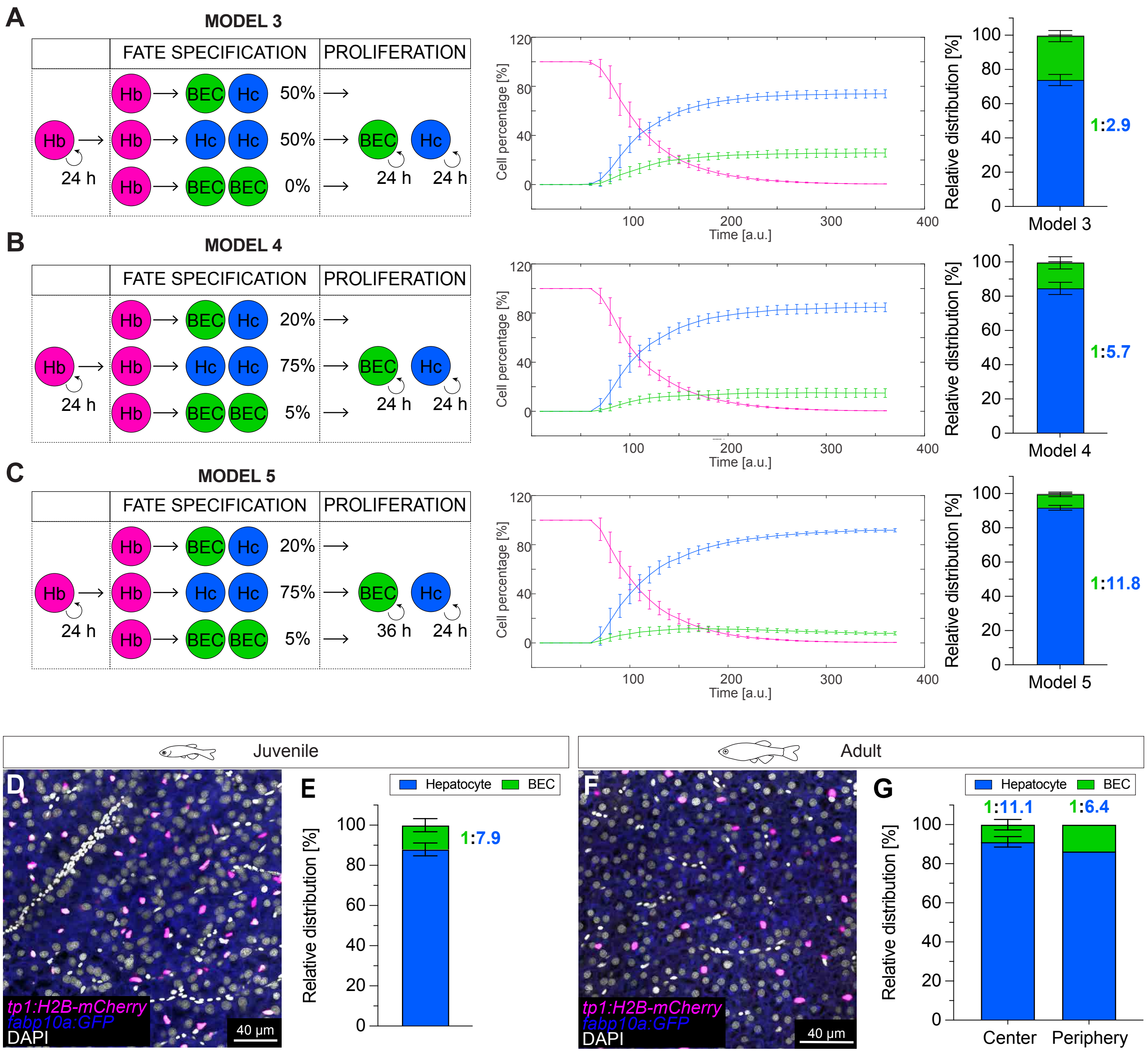


Unterweger et al - Supplementary Figure 1

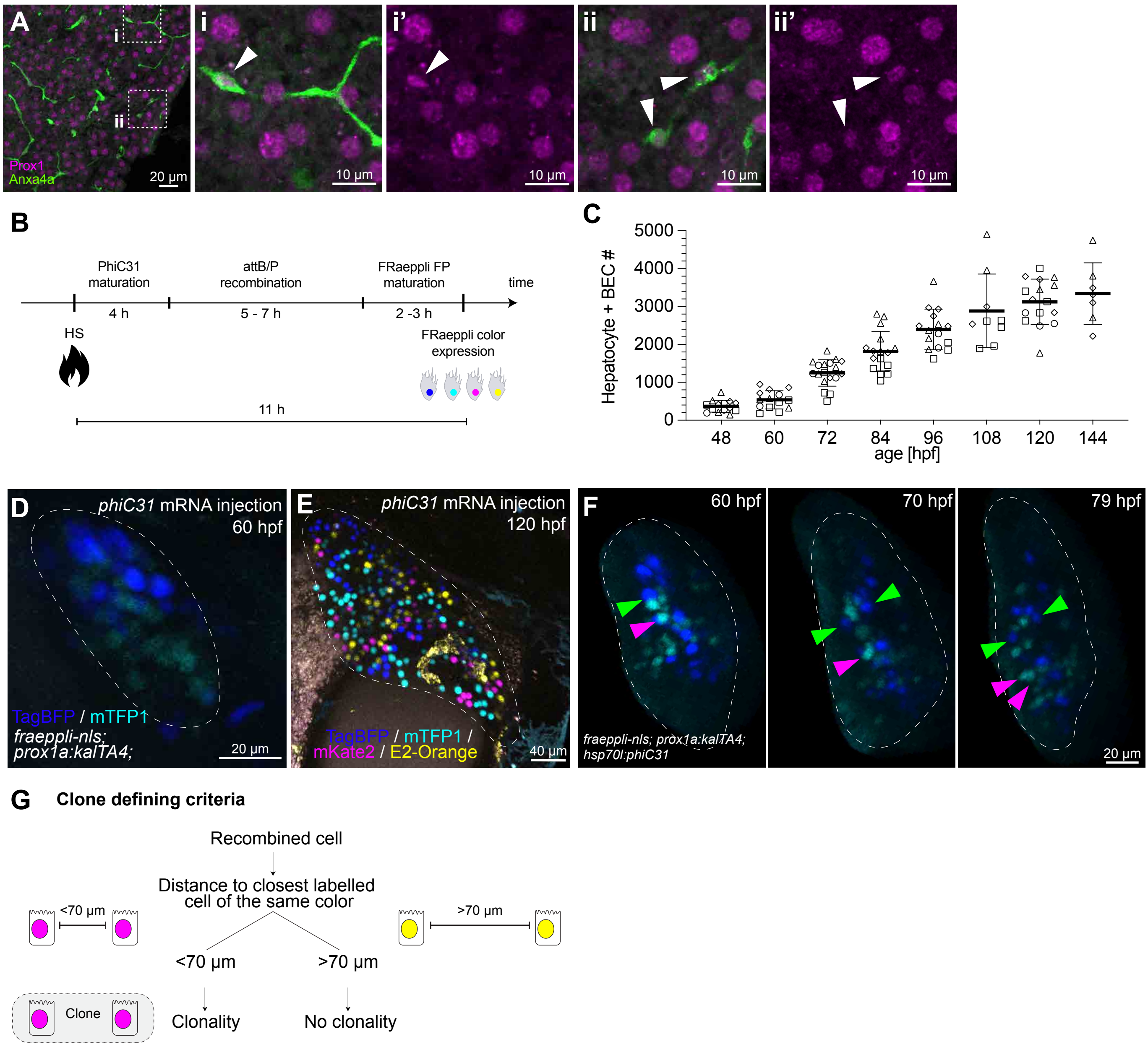


Unterweger et al - Supplementary Figure 2

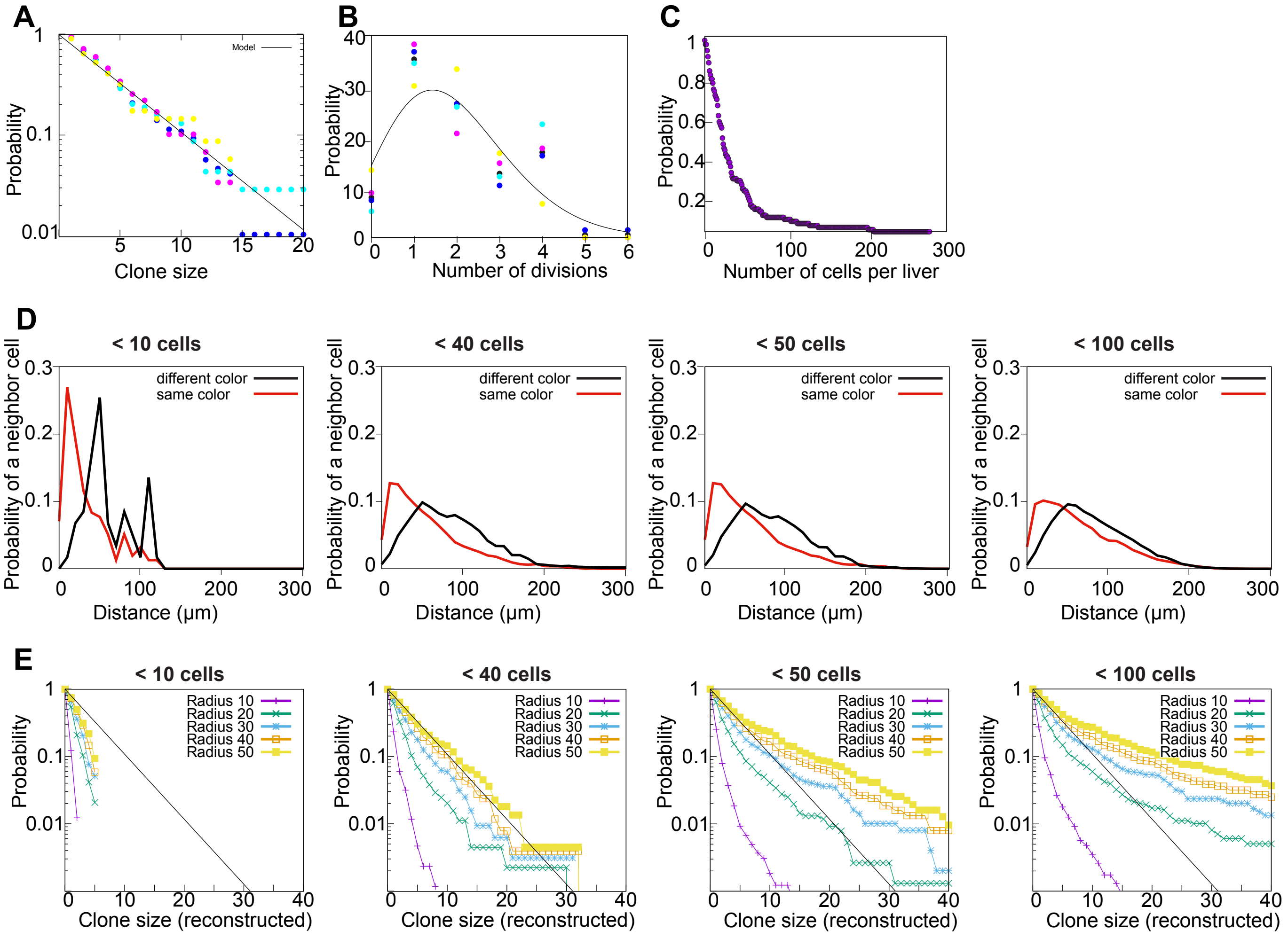


Unterweger et al - Supplementary Figure 3

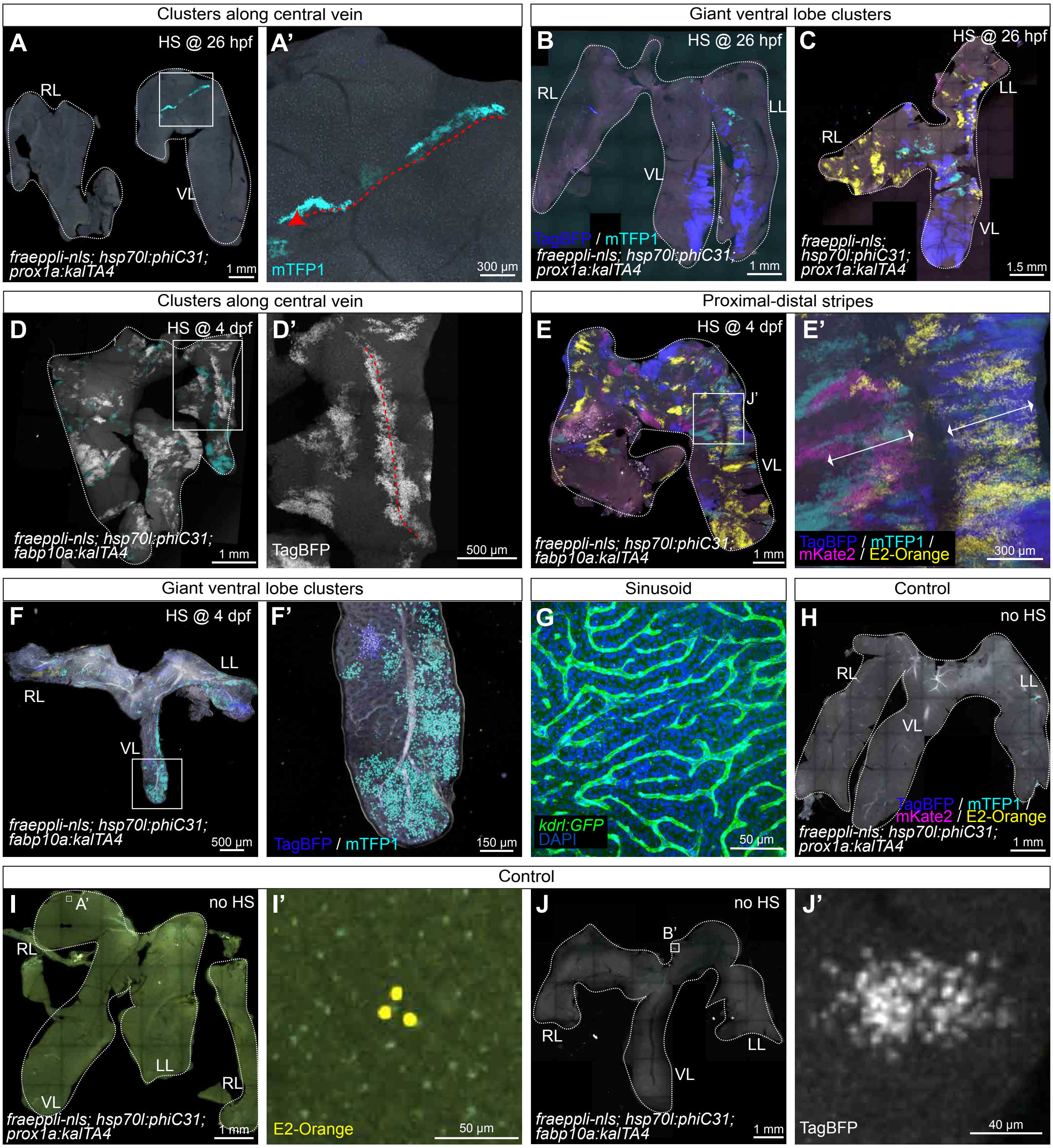


Unterweger et al - Supplementary Figure 4

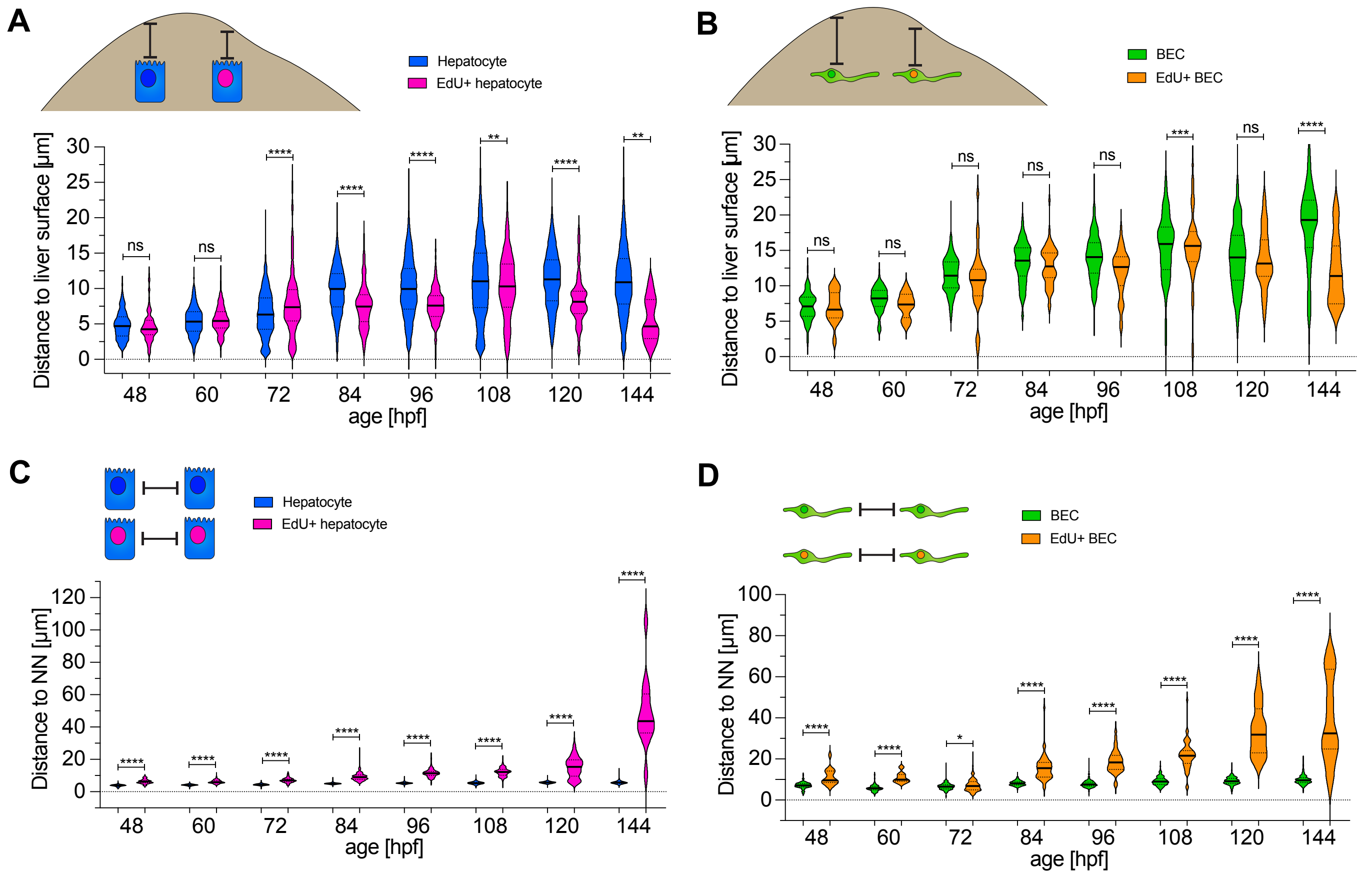


Unterweger et al - Supplementary Figure 5

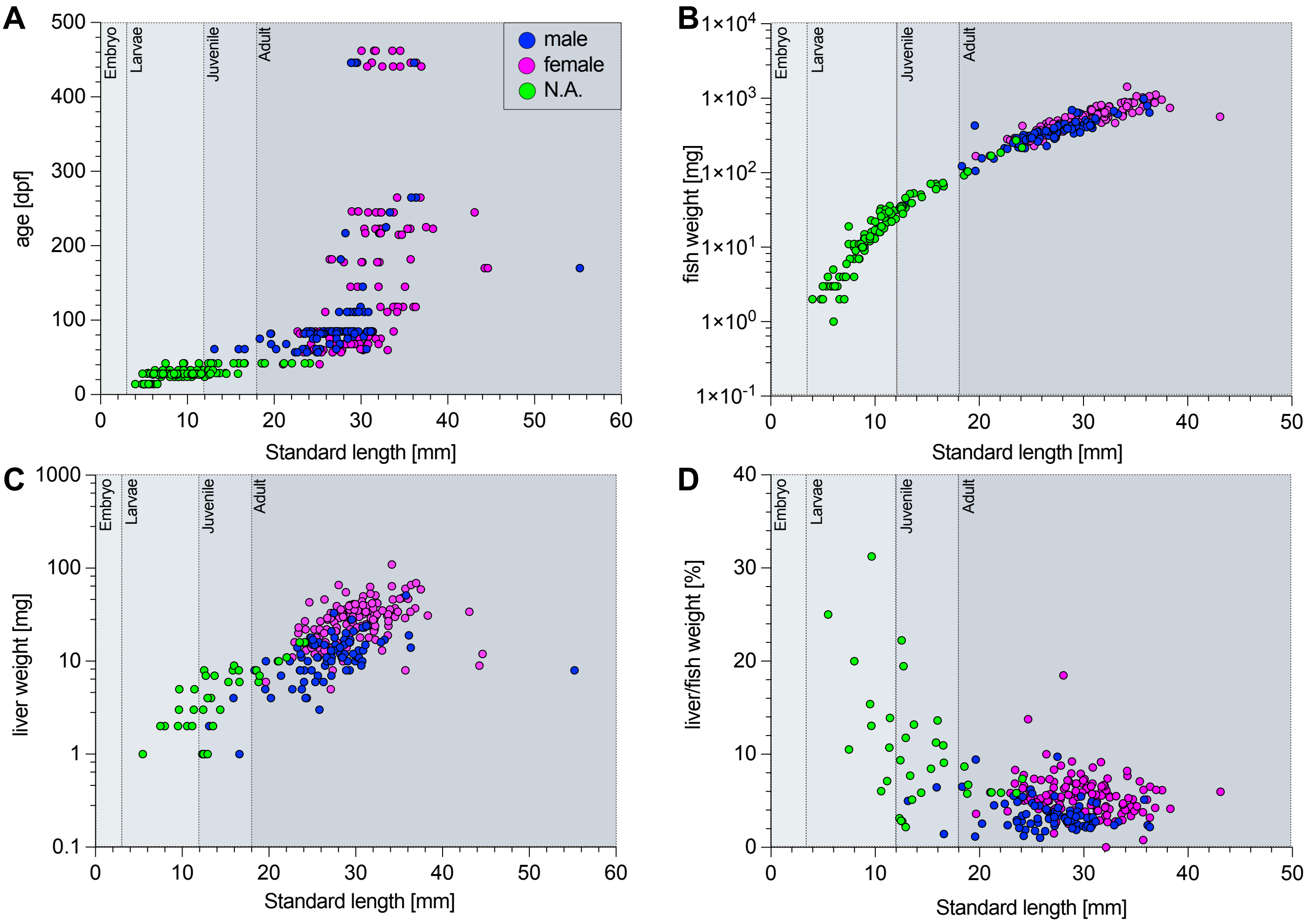


Unterweger et al - Supplementary Figure 6

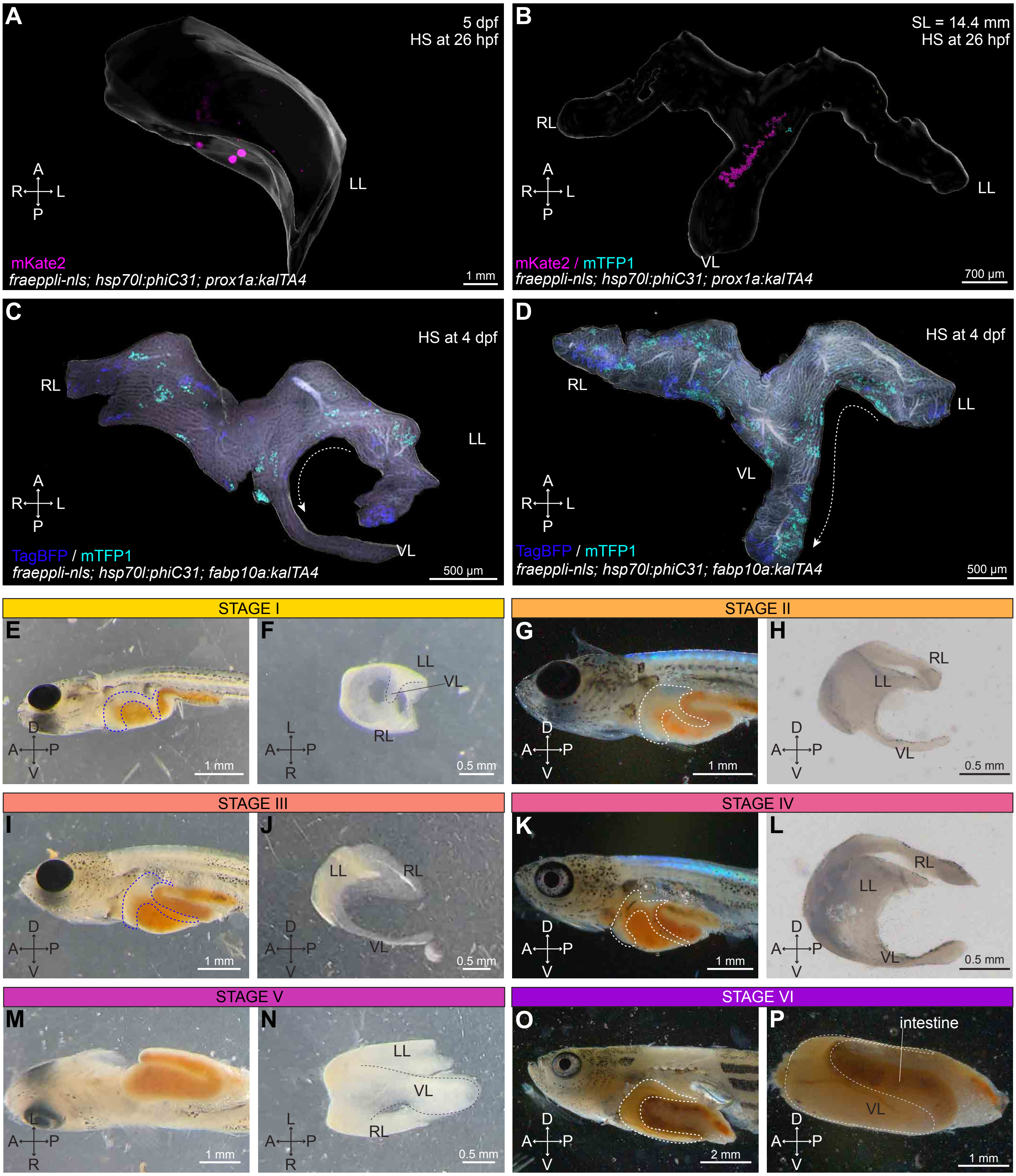


Unterweger et al - Supplementary Figure 7
